# High content live profiling reveals concomitant gain and loss of function pathomechanisms in C9ORF72 amyotrophic lateral sclerosis

**DOI:** 10.1101/2020.04.15.040394

**Authors:** Arun Pal, Benedikt Kretner, Masin Abo-Rady, Hannes Glaß, Marcel Naumann, Julia Japtok, Nicole Kreiter, Tobias M. Böckers, Jared Sterneckert, Andreas Hermann

**Affiliations:** Division of Neurodegenerative Diseases, Department of Neurology, Technische Universität Dresden, Fetscherstr. 74, 01307 Dresden, Germany; Center for Regenerative Therapies TU Dresden (CRTD), Technische Universität Dresden, Fetscherstr. 105, 01307 Dresden, Germany; Translational Neurodegeneration Section „Albrecht-Kossel“, Department of Neurology, and Center for Transdisciplinary Neuroscience (CTNR), University Medical Center Rostock, University of Rostock, Gehlsheimer Str. 20, 18147 Rostock, Germany; German Center for Neurodegenerative Diseases (DZNE) Rostock/Greifswald, Gehlsheimer Straße 20, 18147 Rostock, Germany; Institute for Anatomy and Cell Biology, Ulm University, Ulm, Germany

## Abstract

Intronic hexanucleotide repeat expansions (HREs) in C9ORF72 are the most frequent genetic cause of amyotrophic lateral sclerosis (ALS), a devastating, incurable motoneuron (MN) disease. The mechanism by which HREs trigger pathogenesis remains elusive. The discovery of repeat-associated non-ATG (RAN) translation of dipeptide repeat proteins (DPRs) from HREs along with reduced exonic C9ORF72 expression suggests gain of toxic functions (GOF) through DPRs versus loss of C9ORF72 functions (LOF). Through multiparametric HC live profiling in spinal MNs from induced pluripotent stem cells (iPSCs) and comparison to mutant FUS and TDP43, we show that HRE C9ORF72 caused a distinct, later spatiotemporal appearance of mainly proximal axonal organelle motility deficits concomitant to augmented DNA strand breaks (DSBs), DPRs and apoptosis. We show that both GOF and LOF were necessary to yield the overall C9ORF72 pathology. Finally, C9ORF72 LOF was sufficient – albeit to a smaller extent – to induce proximal axonal trafficking deficits and increased DSBs.

**Single sentence summary:** Pathogenesis in C9ORF72 ALS shows a distinct spatiotemporal axonal organelle trafficking impairment caused by gain and loss of function mechanisms.

## Introduction

ALS is a devastating, incurable motoneuron (MN) disease. Hallmarks of ALS pathology are degeneration of spinal and cortical MNs causing progressive muscular paralysis leading to death within 2-5 years after the onset of clinical manifestation (*1*). MN degeneration progresses by retraction and dying-back of axons from neuromuscular junctions to final death of somata (*2–4*). We have previously modelled retrograde axonal dying-back *in vitro* in a human cell model using compartmentalized iPSC-derived spinal MNs from ALS patients (*5*). A better understanding of the underlying pathomechanism is hampered by the multitude of genetic causes in familial and sporadic ALS. To date, over 30 distinct mutations have been identified (*6, 7*), ranging from single amino residue substitutions to truncations and intronic HREs. This diversity of affected genes and mutation types seems to contradict the common scheme of MN degeneration and final clinical outcome in ALS and calls for a thorough, comprehensive dissection in clinically relevant models to reveal mutation-specific upstream versus more common downstream mechanisms during the progression of neurodegeneration. To this end, we are using fast multichannel live imaging on compartmentalized axons *in vitro* at standardized distal versus proximal readout sites (*5*) owing to the hotly debated role of membrane trafficking defects in many neurodegenerative diseases (*8–10*). Using this setup, we have previously reported about deficient mitochondrial and lysosomal organelle trafficking in iPSC-derived spinal motor neurons from ALS patients bearing mutant FUS (*5*) and TDP43 (*11*), two frequent genetic causes of ALS.

HREs of GGGGCC in intron 1 of C9ORF72 are the most frequent cause of ALS, accounting for 40% of familial and 5% of sporadic cases (*12*). HREs in C9ORF72 is also one of the main genetic causes of frontotemporal dementia (FTD) (*13, 14*). GGGGCC repeat numbers range from 2-23 in healthy persons and are increased to at least sixty in ALS patients and beyond one thousand in extreme cases (*13, 14*). C9ORF72 is inherited in an autosomal dominant fashion and the mechanism by which HREs cause ALS remains unclear. Since HREs concomitantly occur with reduced expression of the exonic C9ORF72 gene (*15–17*), a loss of its reported function (LOF) in axonal trafficking due to haploinsufficiency appeared feasible (*18–20*). However, various knockout (KO) zebrafish and mouse models of C9ORF72 failed to recapitulate MN degeneration and ALS pathology (*21–23*). Subsequently, the discovery of non-canonical RAN-translation of neurotoxic DPRs from intronic HREs (*24, 25*) led to the hypothesis of a DPR-mediated GOF (*22*). Specifically, DPRs are translated bidirectionally from both the sense and antisense HRE-RNA transcripts resulting in a whole spectrum of DPR variants, amongst them more abundant poly glycine-alanine (GA) and poly glycine-proline (GP) (*26, 27*). DPR expression confers its toxic GOF presumably through the formation of inclusion bodies which sequester phosphorylated Ataxia Telangiectasia Mutated (pATM), a key player of DNA damage response, and heterogeneous ribonucleoprotein (hnRNP) A3, normally limiting DPR expression (*26, 27*). The resultant DNA damage accumulation eventually leads to neurodegeneration. Other HRE-mediated GOF mechanisms in concert comprise the formation of sense and antisense RNA repeat-expansion foci (RREs) resulting from bidirectional transcription (*27*). RREs confer RNA toxicity through erratic RNA processing and splicing. Moreover, during the transcription of HREs, nascent RNA is prone to hybridize with the DNA template strand, thereby displacing the complementary DNA strand and forming a three-stranded structure termed R-loops (*27, 28*) which directly increase the risk of DNA strand breaks (DSBs). But again, some mouse models testing GOF by overexpressing HREs and DPRs failed to recapitulate MN degeneration, particularly dying-back events and ALS pathology (*29, 30*) whereas a recent novel transgenic mouse model expressing more toxic poly-PR showed at least some loss of spinal MNs (*31*), suggesting that DPR composition and expression technicalities of mouse models matter.

We have recently established isogenic lines of iPSC-derived spinal MNs comprising parental C9ORF72 from ALS patients along with a (i) gene-corrected (GC) variant with intronic HREs excised, (ii) a KO of the exonic C9ORF72 part with intronic HREs maintained and (iii) a similar KO of C9ORF72 in control cells with naturally no HREs (*32*). To this end, we have utilized here HC phenotypic live profiling of mitochondrial and lysosomal organelle trafficking in axons of HRE C9ORF72 MNs. This approach appeared particularly attractive in light of the documented roles of C9ORF72 in membrane trafficking (*17, 18, 20*). Deficient trafficking in aged C9ORF72 was mirrored by DNA damage and DPR accumulation along with apoptosis. LOF of exonic C9ORF72 in the KO variant was further exacerbating perturbed trafficking and apoptosis due to the remaining HRE-mediated GOF whereas the GC variant with no HREs was fully rescuing all phenotypes. Surprisingly, the ‘pure’ LOF of exonic C9ORF72 in the KO variant of control cells with naturally no HREs partially mimicked the trafficking, DNA damage and apoptosis phenotype, thereby arguing against a sole role of HRE-mediated GOF.

## Results

### Live imaging of compartmentalized MNs revealed distinct organelle trafficking defects in C9ORF72 compared to FUS and TDP43

C9ORF72 has reported roles in endosomal and autophagic membrane trafficking (*17, 18, 20*). Furthermore, MNs with HREs in C9ORF72 showed decreased lysosomal axonal trafficking compared with gene corrected MNs (*32*). Thus, we first wanted to compare trafficking deficits in C9ORF72 lines to other typical ALS causing genes, i.e. FUS and TDP43.

We selected a gender mix of five different ALS patients with confirmed heterozygous HREs in intron 1 of the C9ORF72 gene locus with repeat numbers between 50 – 1800 (C9-1, C9-2, C9-3, C9-4, C9, table 1), respectively, and compared them against three healthy control donors (Ctrl1, Ctrl2, Ctrl3, table 1). These lines were fully characterized and validated in previous publications (table 1). Furthermore, we included our recently published phenotypic profiles from mutant TDP43 and FUS (*33*) to compare them against HRE C9ORF72. All iPSC lines were matured to spinal MN in microfluidic chambers (MFCs), in which only axons could reach and fully penetrate the microgroove barrier of channels from the proximal soma seeding site to distal exits (*5*) (fig. 1a), thereby enabling axon-specific studies with defined antero- versus retrograde orientation. Of note, our differentiation protocol combined with 900µm length of microgrooves resulted in exclusive penetration by MN axons, as we documented previously (*33*). We performed fast dual color live imaging of mitochondria and lysosomes at strictly standardized distal versus proximal readout positions as described (*5*) at day D21. All movies were analyzed with the FIJI TrackMate plugin to deduce organelle tracks with respect to mean speed and track displacement, the latter serving as measure for directed, processive movements as opposed to random walks. Our movie analysis was previously established to reveal distal axonal trafficking defects in mutant TDP43- and FUS-ALS (*5, 11*). Maximum intensity projections of entire movie stacks enabled a preliminary visual inspection for major alterations in motility patterns (Fig. 2a, movie 1, 2). Directed, processive trafficking events were highlighted as long trajectories, whereas stationary organelles and non-processive ‘jitter’ remained virtually as punctae. We obtained HC phenotypic signatures for each line as recently described (*33*). In brief, each parameter was expressed as Z-score deviation from pooled control lines at the proximal readout and plotted with a connecting line to obtain the signature (*33*) (fig. 1b). A master set of 11 parameters was obtained four times owing to two readout positions (distal versus proximal) and two markers (Mito- and Lysotracker), assembled to a signature of 44 parameters in total (*33*). Only Z-scores below −5 and above 5 were considered as significant deviations due to established conventions (*33*) (grey horizontal lines, fig. 1b). As all HRE C9ORF72 lines showed very similar signatures at D21 (in pink, fig. S2) we averaged their data to obtain a pooled profile (C9ORF, in red, fig. 1b). In essence, C9ORF displayed a flat line similar to controls at D21 (red versus blue, fig. 1b), consistent with no phenotype in the raw data (Fig. 2a, S1, movie 1, 2), in track displacement and mean speed (Fig. 2c, movie 1, 2). The FUS (in grey) and TDP43 (in green) signatures (fig. 1b) were virtually identical to our recent report (*33*) except they were normalized to the pooled control lines used in this study (Ctrl, table 1) and not to the isogenic FUS gene-corrected line (*33*). Again, mutant FUS and TDP43 showed pronounced reductions in many parameters, which were particularly pronounced in distal FUS axons, essentially indicating a distal axonopathy in both, TDP43 and FUS. While the Z-scores indicated to what extend a single parameter deviated from control conditions, we strived to have an objective measure of comparing entire multiparametric signatures and grouped them based on similar phenotypic traits. We generated a hierarchical cluster dendrogram with KNIME as described (*33*) (fig. 1c, d). As expected, FUS and TDP43 were each assigned to distinct clusters with FUS being even more deviated, whereas C9ORF and Ctrl cells were grouped together in a cluster termed ‘physiological’ (fig. 1c, d), consistent with no observed cross phenotype at D21 (fig. 1b, 2a, c, S1, S2, movie 1, 2). In summary, hierarchical clustering confirmed the distinct phenotype of mutant FUS and TDP43 confined to the distal axon as opposed to the lack or very small alteration in C9ORF at D21.

**Table 1.**
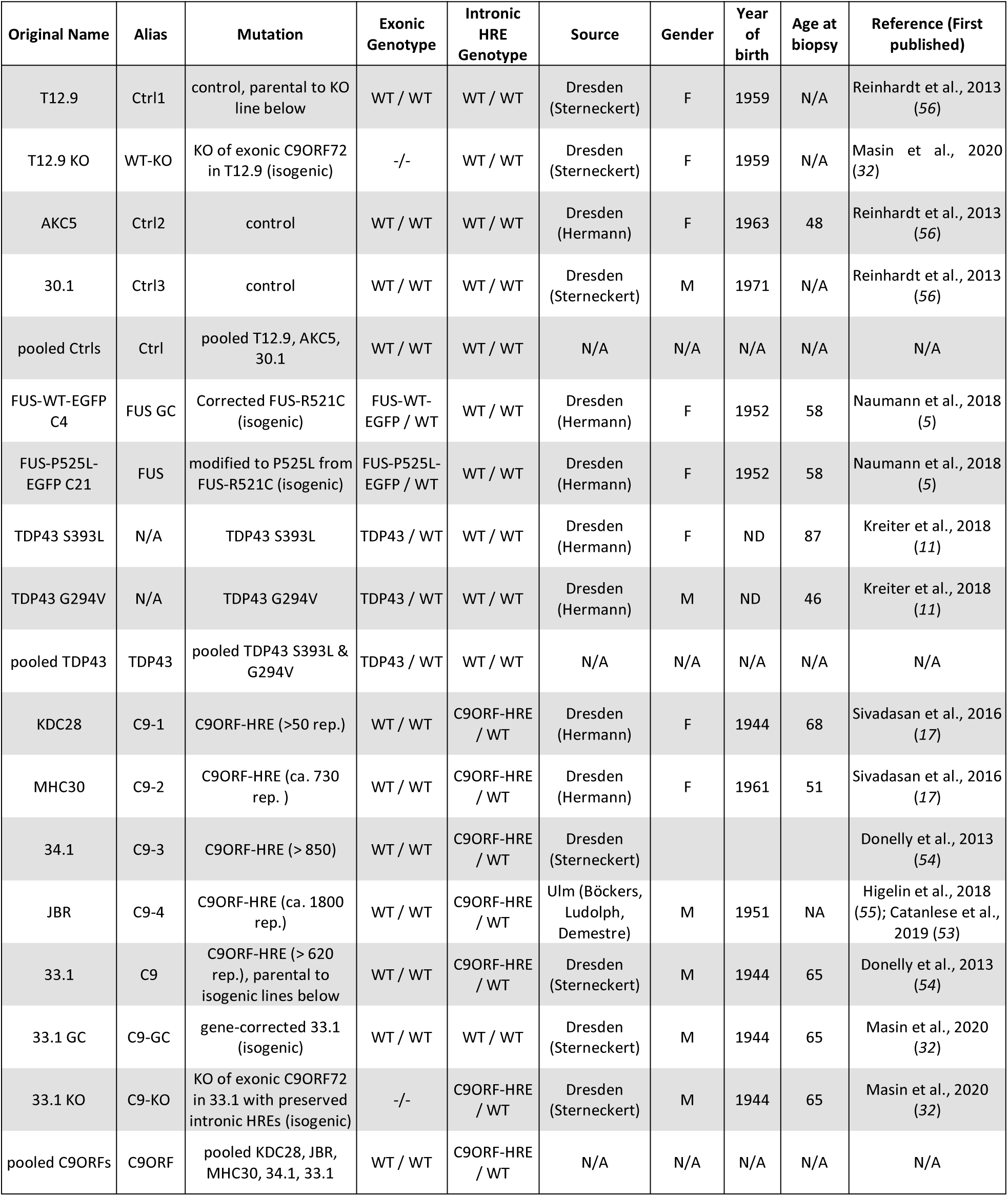
Overview cell line characteristics

**Figure 1.**
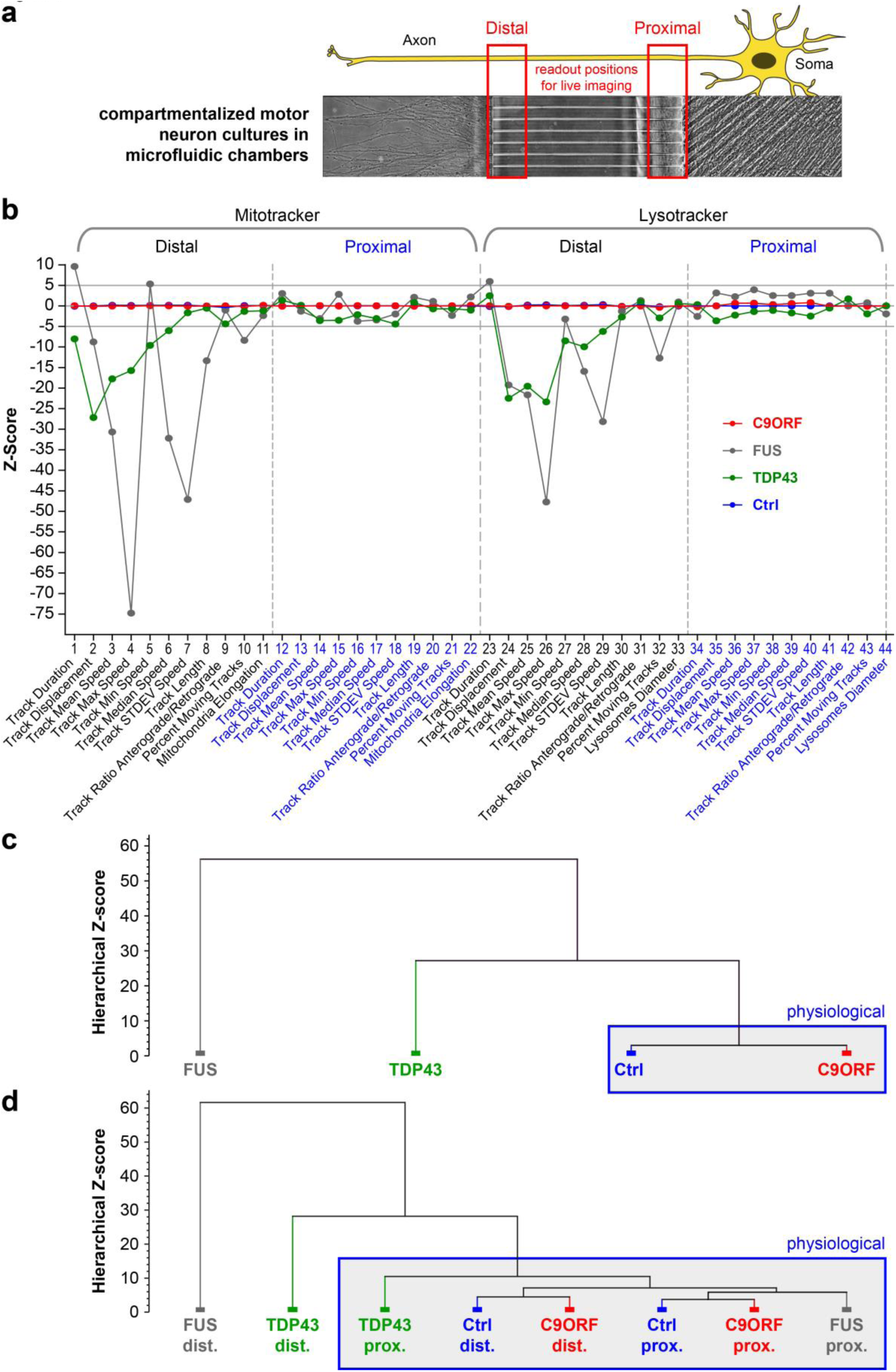
Multiparametric HC phenotypic profiling revealed no phenotype in C9ORF72 at D21 of spinal MN maturation. **(a)** Schematic live set up of motor neurons (MNs) in Zona microfluidic chambers (MFCs). **(b)** The central microgroove of channels formed a physical barrier between the distal (left) and proximal (right) site where the somas were seeded. Only axons, not dendrites, could penetrate the microchannels. **(c)** Multiparametric HC signatures corresponding to the maximum intensity projections in fig. 2a, D21. Shown is the Z-score deviation of each tracking parameter from the proximal readout of pooled Ctrl lines (in dark blue). A set of 11 parameters (bottom labels) was deduced for both the Mito- and Lysotracker, distal versus proximal each, as indicated in the header, resulting in 44 parameters in total. The signatures of mutant FUS (in red), its corresponding isogenic gene-corrected control (FUS GC, in light blue) and TDP43 (in green) were taken from our previous publication to facilitate the comparison to C9ORF (in brown, pooled lines, for individual lines refer to fig. S2). Horizontal grey lines at 5 and −5 indicate significance thresholds. Note the nearly unaltered trafficking in FUS and TDP43 at the proximal readout as opposed to strong negative parameter deviations at the distal site in FUS distinct from the more modest phenotype of TDP43. Conversely, C9ORF exhibited a flat line similar to control lines, consistent with no phenotype at D21 (fig. 2a, c). **(c)** Hierarchical cluster dendrogram of entire signatures shown in (a). The hierarchical Z- score (ordinate) indicates the deviation of entire signatures from each other and is not to be mistaken with the individual parameter Z-scores in (b). Blue boxed cluster highlights physiological signatures. Note how the phenotypically unremarkable C9ORF (brown) clustered with Ctrl (dark blue) against the deviate TDP43 (green) and more severe FUS mutant (red). **(d)** Hierarchical cluster dendogram of partial signatures comprising either only all distal or proximal parameters (Mitotracker and Lysotracker, respectively) to compare site – specific phenotypes. Note how both Ctrl parts (dark blue, distal and proximal) clustered closely with the proximal FUS part (red) on the right into a physiological super cluster boxed in blue due to the close physiological trafficking state whereas the drastic organelle arrest in the distal FUS part on the far left (red) was highly distinct to its physiological parts at the proximal site. TDP43 showed some moderate deviation in its proximal part (green) within the physiological super cluster and a clear deviation in its distal part (green), albeit less drastic than FUS (red). C9ORF was contained in the physiological super cluster with both the distal and proximal part due to no phenotype at D21.

**Figure 2.**
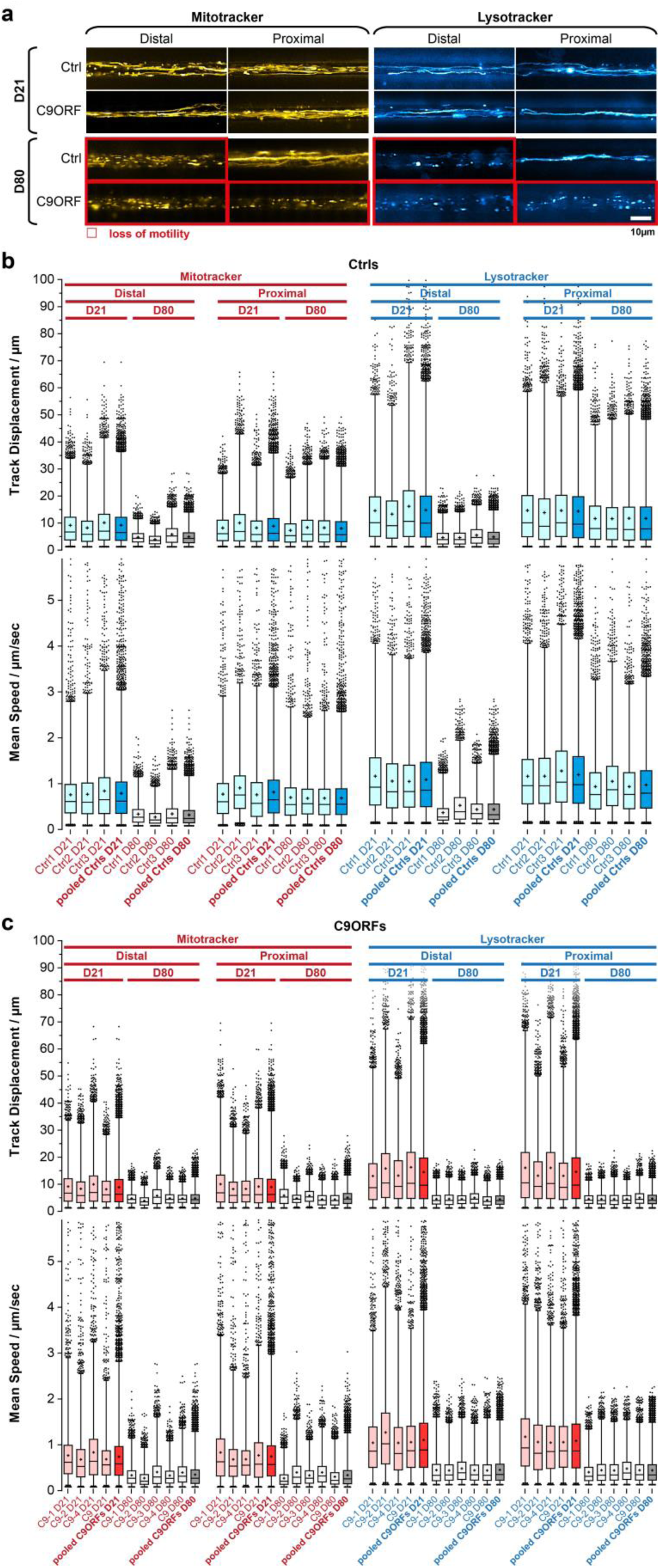
Loss of organelle motility in aged C9ORF72 spinal MNs. **(a)** Maximum intensity projections of movie raw data acquired live with Mitotracker (left) and Lysotracker (right) at the distal (left) versus the proximal (right) microchannel readout position as shown in panel a. Movies were acquired at 21 days during maturation (top galleries, D21) versus aged stage at D80 (bottom galleries). Red boxed images highlight loss of motility. Note the loss of motility at D80 in both distal and proximal C9ORF axons as opposed to distal loss only in Ctrl neurons. Shown are the Ctrl2 and C9-1 line (table 1) as representative example. An overview of all lines is provided in fig. S1. Scale bar = 10µm. **(b)** Organelle tracking analysis corresponding to (a) of all Ctrl lines as box plots, Mitotracker (left) and Lysotracker (right) distal versus proximal and D21 versus D80 as indicated in the header. Whiskers: 1-99%, box: 50%, horizontal line: median, cross: mean, outliers: black dots. Shown are the individual Ctrl lines (Ctrl1-3, in pale blue or light grey) along with the pooled analysis (Ctrls pooled, in full blue or dark grey), as indicated in bottom labels. The tracking analysis was performed for organelle track displacement (top box plots) and mean speed (bottom box plots). Physiological motility is indicated in pale/full blue as opposed to organelle arrest in light/dark grey. Note that proximal motility remained physiological over ageing (D21 and D80) in all control lines as opposed to the distal decline at D80. **(c)** Same as (b) but for all C9ORF lines. Shown are the individual lines (C9-1, C9-2, C9- 3, C9-4, C9 in pale red or light grey) along with the pooled analysis (C9ORFs pooled, full red or dark grey), as indicated in bottom labels. Physiological motility is indicated in pale/full red as opposed to organelle arrest in light/dark grey. Note that both distal and proximal motility was lost over ageing (compare D21 and D80) in all C9ORF lines whereas proximal control organelles in (c) remained motile over ageing. C9-3 was not measured at D21.

### Organelle trafficking defects deteriorate during ageing in HRE C9ORF72

Different to FUS-ALS, C9ORF72-ALS is a classical old onset form of ALS (*34*). Thus, we hypothesized that degenerative phenotypes will be visible at later time points and performed fast dual color live imaging of mitochondria and lysosomes at D21 and D80. All HRE C9ORF72 lines exhibited normal, physiological distal and proximal organelle motility at D21 similar to control cells (Fig. 2a, S1, movie 1, 2). After ageing until D80, control cells apparently lost their processive organelle motility at the distal site (Fig. 2a, S1, movie 1, 2, boxed in red), whereas their proximal trafficking appeared unaltered, feasibly due to a “physiological” retrograde dying back of axons over extended ageing (*5*). Conversely, all HRE C9ORF72 lines exhibited a decline in trafficking at D80 at both the distal and proximal site, suggesting a distinct progression of neurodegeneration (Fig. 2a, S1, movie 1, 2, boxed in red).

For each organelle type, we displayed mean speed and track displacement resulting from the tracking analysis as box plots (fig. 2b, c). All control (fig. 2b) and HRE C9ORF72 (fig. 2c) lines exhibited, in essence, two rather discrete trafficking states: one mobile (blue boxes for all control lines, fig. 2b; red boxes for all HRE C9ORF72 lines, fig. 2c) with an average track displacement around 9/14µm (Mito-/Lysotracker) and mean speed of about 0.7/1.1 µm/sec (Mito-/Lysotracker) as opposed to one relatively immobile (grey boxes, fig. 2b, c) with an average track displacement around 5/5µm (Mito-/Lysotracker) and mean speed of 0.4/0.4 µm/sec (Mito-/Lysotracker). At D21, all lines (control and HRE C9ORF72) displayed the mobile state at either site (distal and proximal) for either type of organelle (mitochondria and lysosomes). At D80, all control lines had their trafficking still fully maintained in the mobile state at the proximal axon site but deteriorated to the immobile state at the distal site (fig. 2b), whereas all HRE C9ORF72 lines showed the immobile state at both the distal and proximal site (fig. 2c), consistent with the apparent pathological decline in the raw data (Fig. 2a, S1, movie 1, 2).

### Multiparametric spatiotemporal HC organelle tracking analysis revealed a proximal axonopathy in HRE C9ORF72

In light of the distal axonopathy in mutant FUS and TDP43 already at D21 along with premature axonal dying back (*5*) and a similar figure in physiological control cells at D80 (fig. 2a, b, S1, S2, movie 1, 2) we envisioned two distinct plausible scenarios to explain the virtual organelle arrest at both the distal and proximal site in C9ORF72 over extended ageing at D80 (fig. 2a, c, S1, S2, movie 1, 2): either dying-back of axons progresses earlier and faster between D21 and D80, thereby reaching to the proximal readout until D80 whereas in control cells the slower dying-back has still no impact here, or the proximal phenotype occurs independently of axonal dying-back by a distinct mechanism.

To understand these spatiotemporal differences in more detail, we analyzed the multiparametric HC signatures at multiple time points over a course from D14-80 (Fig. 3a, S2). Given the high similarities within Ctrl and C9ORF lines, respectively, fig. 3 shows pooled signatures (Ctrl: dark blue; C9ORF: red) whereas fig. S2 provides the corresponding profiles of individual lines. At D14, 21, and 28 Ctrl and C9ORF lines displayed virtually flat lines as signatures, consistent with no cross phenotype until D28 (fig. 3a). However, from D40 onward we observed negative deviations in distal control axons (dark blue profiles, fig. 3a) for most speed parameters, track duration and displacement for either type of organelle. The overall reduction of organelle motility further exacerbated but remained restricted to the distal site in controls until D80 (dark blue profiles, fig. 3a). Only few proximal lysosome parameters finally showed borderline significance in their alterations that were, however, marginal compared to the drastic impairments at the distal site.

**Figure 3.**
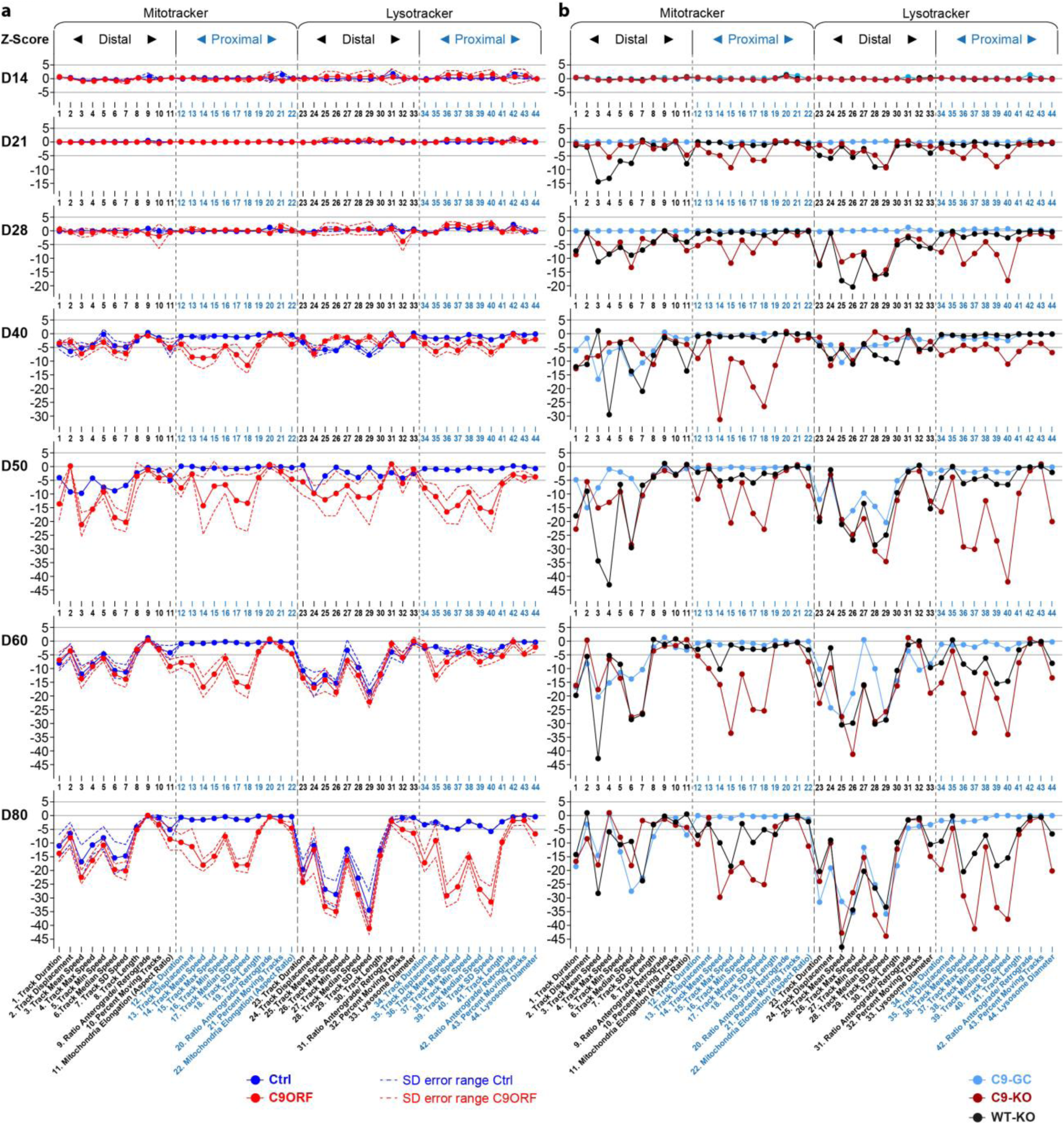
HC phenotypic profiling over extended time course revealed global axonal trafficking defects in C9ORF72 spinal MNs over ageing. **(a)** Multiparametric HC profiles of pooled Ctrl (in blue) and C9ORF lines (in red) were deduced as in fig. 1b over a time course from D14 – D80 as indicated on the left. For overview of all individual lines refer to fig. S2. Z-scores for all time points were calculated with respect to pooled control lines at D21 proximal. Dotted lines indicate error ranges (standard deviation between lines). Note the onset of trafficking decline (negative parameter deviations, Z-scores ≤ −5) for either organelle type (Mito – and Lysotracker) from D40 onwards that progressed only at the distal readout site in control lines as opposed to the global phenotype in C9ORF72, i.e. simultaneous emergence at both the distal and proximal site. **(b)** Similar time course for (i) C9-GC with excised intronic HREs (in light blue), (ii) C9-KO with intronic HREs preserved but with KO of the exonic C9ORF72 part (in brown), and (iii) WT-KO cells with the same exonic KO and naturally having no intronic HREs (in black). Note how the proximal trafficking decline in C9-GC was restored to physiological levels over the entire time course while the distal decline remained unaltered (compare light blue profiles with dark blue counterparts in (a)). As for C9-KO, note the earlier onset of global (distal & proximal) trafficking defects already at D21 (Z-scores ≤ −5) compared to D40 in parental C9 (compare brown profiles with red counterparts in (a)). As for WT-KO, note the earlier onset of distal trafficking defects already at D21 and the emergence of a proximal decline at later time points as well (from D50 onwards) as opposed to distal decline only in parental Ctrl (compare black profiles with dark blue counterparts in (a)).

In contrast, C9ORF cells exhibited such trafficking defects always simultaneously at both the distal and proximal site with an onset at D40 (red profiles, fig. 3a). The further progression of these defects appeared unsteady in some signatures parts. Particularly for proximal lysosomes, we observed a rapid parameter decrease from D40 to 50, followed by some transient stagnation at D60 towards a most severe impairment at D80 (red profiles, fig. 3a). The concurrent emergence of trafficking defects at both the distal and proximal site from D40 onwards was a surprising finding arguing against a simple retrograde dying-back of the axon but pointing towards a proximal axonopathy by a distinct mechanism.

### Both gain and loss of function contribute to trafficking deficiency in C9ORF72

There is extensive discussion about the pathomechanisms of C9ORF72 HRE including mainly gain of toxic functions (GOF) (*26, 27*) versus loss of C9ORF72 functions (LOF) (*18, 20*). Thus, we wished to investigate the role of GOF and LOF mechanisms in HRE-mediated trafficking defects. To this end, we used CRISPR-Cas9n-mediated gene editing on the parental HRE line C9 (table 1) to generate an isogenic set of additional MN models. A full description and characterization of these KO and GC variants was recently published (*32*) (table 1), including a verification of eliminated C9ORF72 expression in all KO variants and of DPRs in the GC variant. We deduced multiparametric HC signatures for the isogenic C9-GC, C9-KO and WT-KO lines (table 1) over the same time course from D14-80 (fig. 3b).

We first analyzed a gene-corrected control line (C9-GC) with excision of the intronic HREs by replacing the HRE in intron 1 with its wild type counterpart of only 1-3 repeats. This led to a full restoration of all proximal parameter deviations for either type of organelle at any time, thereby resulting in wild-type like signatures with a similar onset at D40 and further progression of distal trafficking defects until D80 without proximal phenotypes (compare light blue profiles in fig. 3b with dark blue in fig. 3a). Hence, gene-correction indicated that all phenotypic perturbations over extended ageing in our C9ORF lines were truly due to C9ORF72 HRE mutation.

Since it is known that HRE-expressing C9ORF72 causes reduced expression of C9ORF72 (*15–17*), we cannot distinguish from these results whether GOF or LOF caused the trafficking phenotypes. To address this, we abolished C9ORF72 expression by excision of the translation start codon in exon 2 as effectively as a conventional gene KO except that HRE-mediated DPR expression from intron 1 still occurred to study their GOF in the absence of C9ORF72 (C9-KO). KO of exonic C9ORF72 while maintaining HREs (C9-KO) led to a premature onset of negative parameter deviations already at D21 (compare brown profiles in fig. 3b with red in fig. 3a), thereby exacerbating the natural C9ORF phenotype.

Having established that KO of exonic C9ORF72 is exaggerating C9ORF72 HRE phenotypes, we aksed whether KO of exonic C9ORF72 in control cells (WT-KO) without HREs is sufficient to cause any phenotypes. By utilizing the same KO method on a healthy control line (Ctrl1) we abolished C9ORF72 expression in a wild type with naturally no HREs to study C9ORF72 LOF in the absence of RAN-translated DPRs (WT-KO). This led to a premature onset at D21 as well, but initially only at the distal site (compare black profiles in fig. 3b with dark blue in fig. 3a). Remarkably, we observed a delayed onset of proximal impairments as well from D50 onwards, finally resulting in a signature at D80 resembling the natural C9ORF profile (compare black profiles in fig. 3b with red in fig. 3a). This peculiar spatiotemporal appearance of this axonal phenotype might suggest that the dying-back of axons progresses earlier and faster between D21 and D80, thereby reaching to the proximal readout until D80 whereas in control cells the slower dying-back has still no impact here. In summary, these results indicate that both GOF of HRE/DPRs and LOF of C9ORF72 protein mechanisms contribute through combinatorial action to the overall C9ORF phenotype.

Since altered onsets and site-specificities (i.e. distal versus proximal axon sites) were emerging as major phenotypic perturbations, we sought to extract these distinctive features from our multiparametric data sets at better clarity. To this end, we summed up the absolute values of all distal versus proximal parameter Z-scores (for both the Mito- and Lysotracker, respectively) to obtain a site-specific measure of overall phenotypic strength relative to control baseline along with the total phenotypic strength of the entire signature (fig. 4, S3). Clearly, Ctrl cells did hardly show any increase of their proximal phenotypic strength over ageing and remained on the initial base level (in blue, top gallery, fig. 4, S3), consistent with no visible phenotype (Fig. 2a, S1, movie 1, 2) and no appreciable proximal parameter deviations in the corresponding signatures (Fig. 3a, S2). By contrast, the distal phenotypic strength was steadily rising from a flat line in Ctrl cells from D40 onwards (in red, top gallery, fig. 4, S3), thereby clearly marking the onset of distal trafficking impairments. C9ORF cells exhibited a rise at both the distal and proximal site (compare blue with red, bottom gallery, fig. 4, S3) from D40 onwards (red profiles, fig. 3a). Excision of intronic HREs (C9-GC) resulted in a flat line at the proximal site over the entire time course indistinguishable from Ctrl cells, consistent with full restoration due to the gene correction (in blue, bottom gallery, fig. 4). Conversely, KO of exonic C9ORF72 with intronic HREs preserved (C9-KO) led to an earlier rise in phenotypic strength from D21 onwards simultaneously at both axon sites (compare blue with red, bottom gallery, fig. 4, S3). Finally, KO of exonic C9ORF72 in control cells with naturally no HREs (WT-KO) resulted in a premature rise at the distal site whereas the proximal rise was delayed to D50 (compare blue with red, top gallery, fig. 4, S3). In summary, the plotting of site-specific phenotypic strength over ageing confirmed the differences in the spatiotemporal progression of axonal trafficking defects at high clarity.

**Figure 4.**
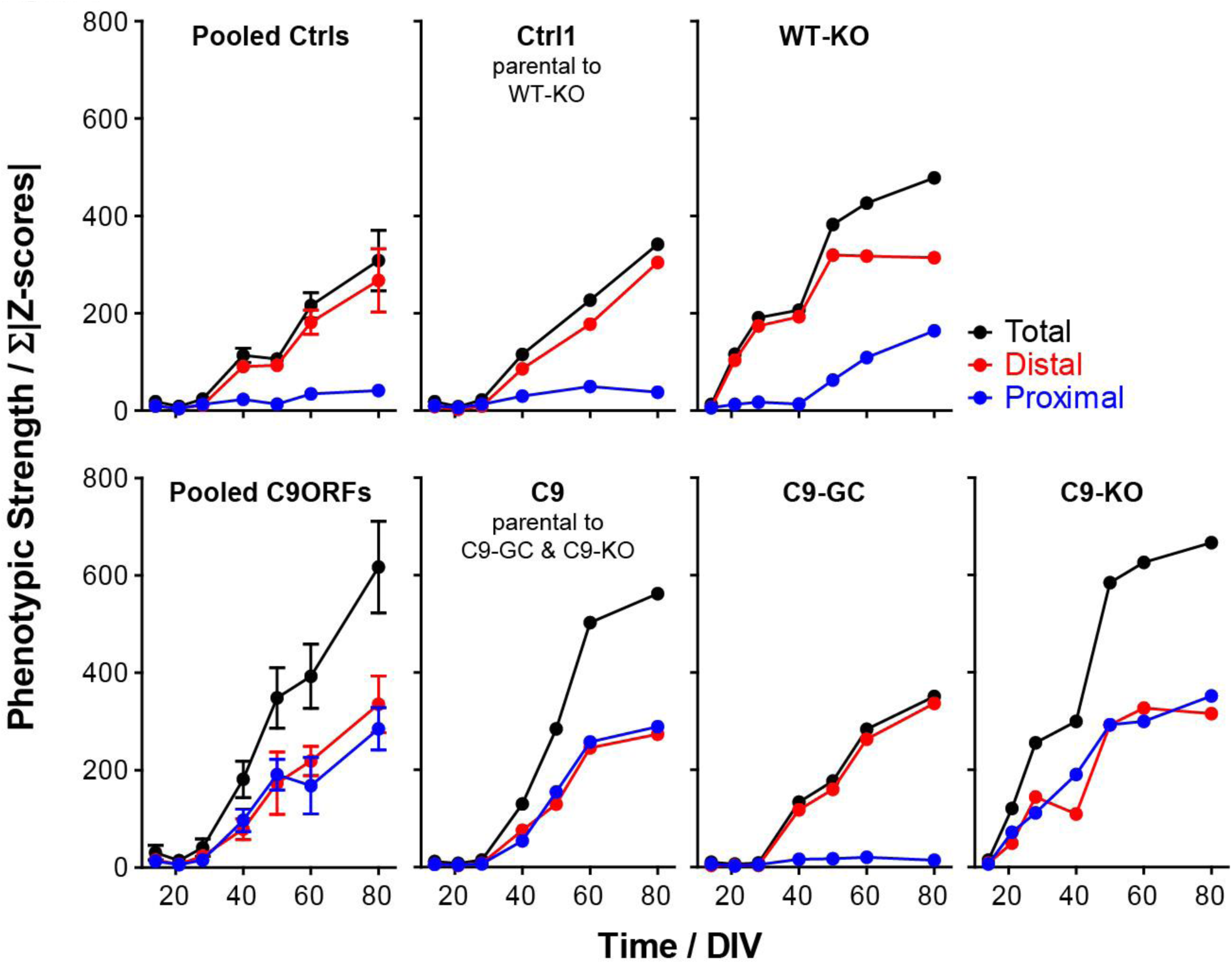
Phenotypic strength progressing over time. The sum of all Z-score moduli of profiles in fig. 3 was plotted as measure of phenotypic strength over time to highlight the progression of declining trafficking. Black curves: entire profile, blue curves: only proximal profile parts, red curves: only distal profile parts. Note how unaltered control cells remained nearly constant on the initial baseline level over the entire time course at the proximal site due to no phenotype (top gallery, Pooled Ctrls and Ctrl1 parental to WT-KO in blue) whereas their phenotypic strength at the distal site exhibited a sudden rise from D40 onwards due to the onset of trafficking decline (top gallery, Pooled Ctrls and Ctrl1 in red). Conversely, WT-KO exhibited a proximal trafficking decline from D50 onwards along with an earlier onset of distal phenotypes already at D21 as compared to D40 in parental Ctrl1 (top gallery). As for parental C9, phenotypic strength exhibited a sudden rise from D40 onwards simultaneously at both the distal and proximal site due to the onset of a global trafficking decline (bottom gallery, Pooled C9ORFs and C9). C9-GC showed a complete absence of the proximal rise in phenotypic strength due to restoration of trafficking at this site while the distal decline remained as in Ctrl cells (bottom gallery, C9-GC). Conversely, C9-KO exhbited an earlier onset of rising phenotypic strength simultaneously at both the distal and proximal site already at D21 as opposed to D40 in parental C9 (bottom gallery, compare C9-KO with C9).

### HRE-mediated axonal organelle trafficking defects concurred with DPR and DNA damage accumulation along with apoptosis

Two commonly recognized hallmarks in the pathology of neurodegeneration are nuclear DNA damage accumulation and apoptosis (*5, 35*). Specifically in C9ORF ALS, HRE- based pathology is believed to be mediated through RAN-translated DPRs (*22, 29, 30*). Thus, we sought to score directly for them with respect to DNA damage and apoptosis when axonal trafficking defects emerged. To this end, we performed immunofluorescence confocal microscopy on the isogenic lines at the end of our time course at D80 versus D21 (i.e. before and after emergence of trafficking defects, fig. 3) to reveal the prominent DPR variants poly glycine-proline (GP) and poly glycine-alanine (*26*) along with the DNA strand break (DSB) markers phospho histone H2A.X (γH2AX, fig. 5, S4, S6) and tumor suppressor 53-binding protein 1 (53BP1, fig. 6, S5, S6) as well as the apoptosis marker cleaved caspase 3 (Casp3, fig. 7) as described (*5*). As for other DPR variants (e.g. GR), we failed to obtained specific staining patterns with available antibodies and rejected them from this study (data not shown). Control cells did only show traces of GP, GA and either DSB marker at both time points (Ctrl1, fig. 5, 6). By contrast, parental C9ORF cells and their KO counterpart exhibited a drastic accumulation of both DPR variants at D80 (C9, C9-KO, fig. 5, 6) whereas at D21 they were indistinguishable from control cells (Ctrl1, fig. 5c, 6b, S4, S5). Specifically for GP, the accumulation occurred as aligned neuritic foci in MAP2-positive neurons (Fig. 5a, green arrowheads). Conversely for GA, the accumulation occurred as larger perinuclear foci (Fig. 6a, green arrowheads), consistent with histological brain sections in C9ORF72 patients in a recent report (*26*). Accumulation of both DPR variants concurred with augmented DSBs, (fig. 5a, 6a, white arrowheads). In case of GP, DSBs were revealed with _γ_H2AX and in case of GA with 53BP1 antibodies due to a species conflict in the co-staining cocktail (see Material and Methods). However, we verified that both DSB markers revealed very similar staining patterns in all lines (fig. S6a, yellow arrowheads) with high colocalization when DSBs augmented (fig. S6b, c).

**Figure 5.**
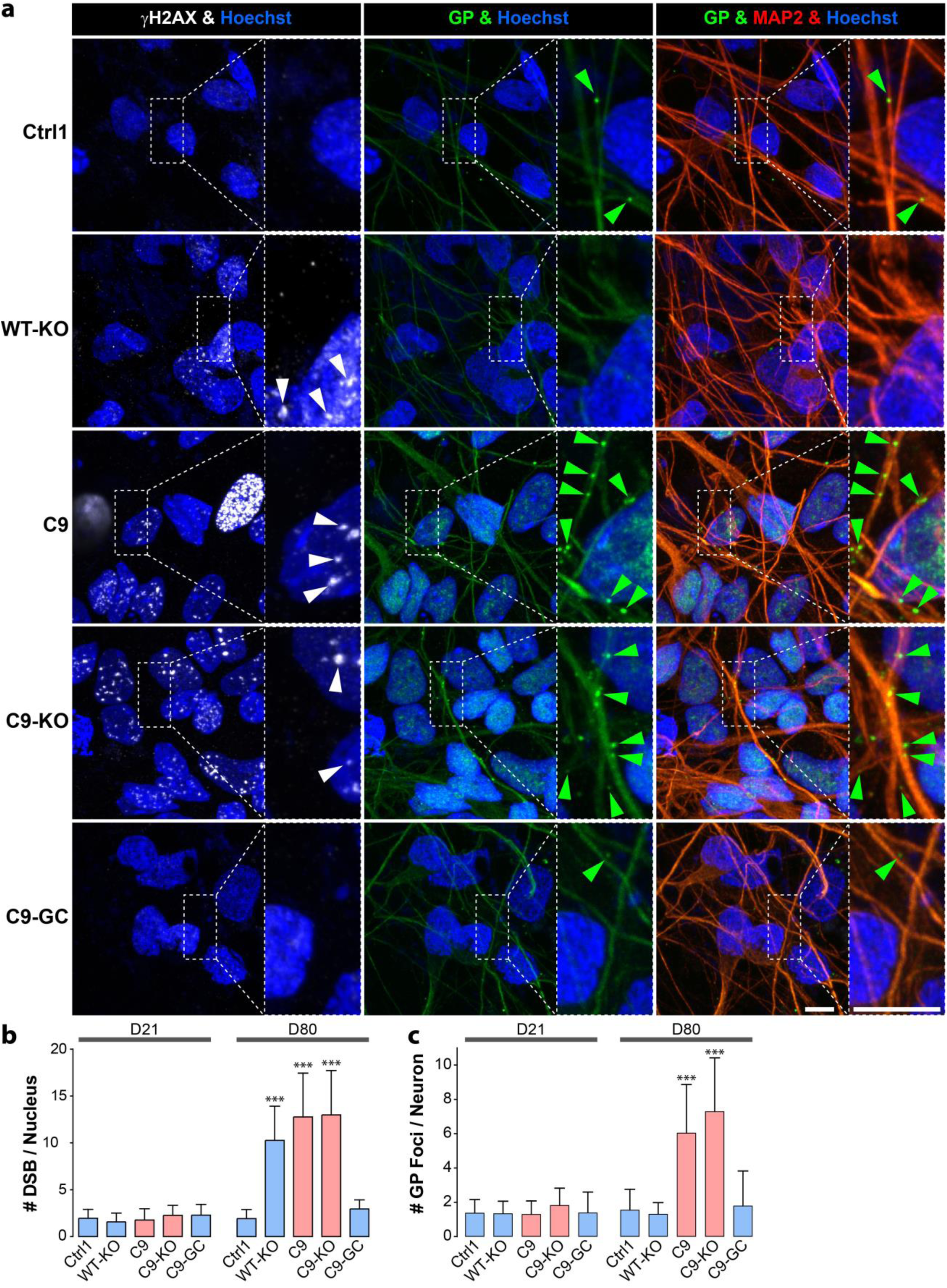
DNA damage accumulation concurred with neuritic GP DPR foci in C9ORF72 spinal MNs over ageing. **(a)** DNA strand break (DSB) marker γH2AX (in white) in Hoechst-positive nuclei (in blue) and neuritic GP foci (in green) in MAP2-positive neurons (in red) were revealed by confocal IF microscopy at D80 endpoints (fig. 3). Dotted boxed areas in image galleries are shown magnified on the right. Note the striking nuclear accumulation of γH2AX- positive nuclear foci (white arrowheads) in parental C9 and C9-KO that was phenocopied by WT-KO. Furthermore, GP foci aligned to neurites (green arrowheads) concurred with nuclear γH2AX accumulation in C9 and C9-KO. Conversely, DSBs and GP foci were nearly absent in C9-GC and parental Ctrl1. Scale bars = 10µm. **(b)** Quantification of (a), number of DSBs (i.e. γH2AX-positive nuclear foci) in MAP2- positive cells at D21 vs D80. For images at D21 refer to fig. S4. Note nearly absent DSBs at D21 in all lines vs drastic DSB accumulation at D80 in parental C9, C9-KO and WT-KO. **(c)** Quantification of (a), number of neuritic GP foci in MAP2-positive neurons at D21 vs D80. For images at D21 refer to fig. S4. Note nearly absent GP foci in all lines at D21 vs aligned foci at D80 in parental C9 and C9-KO. **(b, c)** Asterisks: highly significant increase in any pairwise comparison with unlabeled conditions, one-way ANOVA with Bonferroni post test, *P≤0.05, **P≤0.01, ***P≤0.001, N=60 images from 3 independent experiments, error bars=SD. All unlabeled conditions (i.e. with no asterisk) were not significantly different amongst themselves in any pairwise comparison.

**Figure 6.**
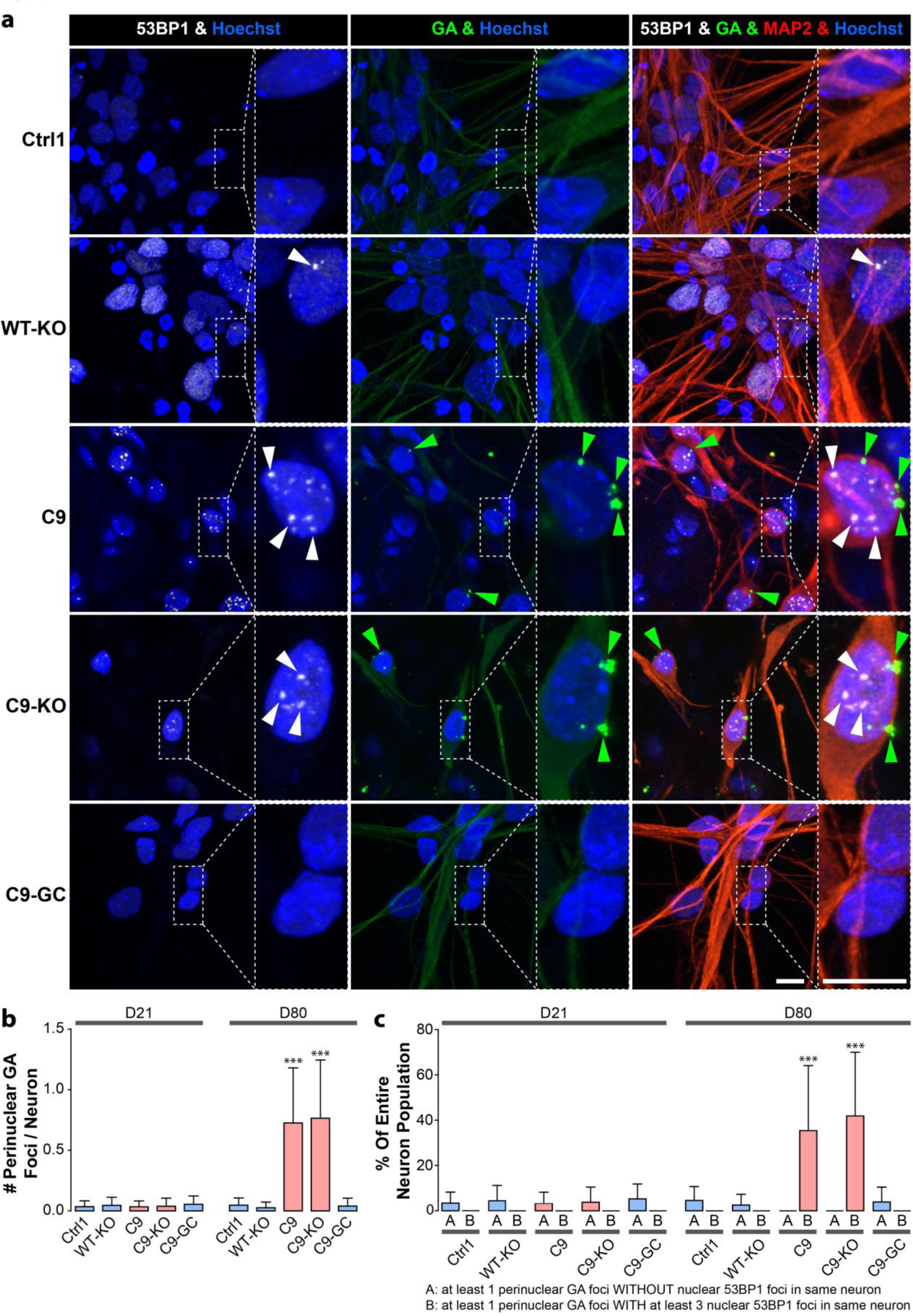
DNA damage accumulation concurred with perinuclear GA DPR foci in C9ORF72 spinal MNs over ageing. **(a)** DSB marker 53BP1 (in white) in Hoechst-positive nuclei (in blue) and perinuclear GA foci (in green) in MAP2-positive neurons (in red) were revealed by confocal IF microscopy at D80 endpoints (fig. 3). Dotted boxed areas in image galleries are shown magnified on the right. Note the striking nuclear accumulation of 53BP1-positive nuclear foci (white arrowheads) in parental C9 and C9-KO that was phenocopied by WT-KO. Furthermore, perinuclear GA foci (green arrowheads) concurred with nuclear 53BP1 accumulation within the same neuron in C9 and C9-KO. Conversely, DSBs and GA foci were nearly absent in C9-GC and parental Ctrl1. Scale bars = 10µm. **(b)** Quantification of (a), number of perinuclear GA foci in MAP2-positive neurons at D21 vs D80. For images at D21 refer to fig. S5. Note nearly absent GA foci at D21 in all lines vs drastic GA accumulation at D80 in parental C9 and C9-KO. **(c)** Quantification of (a), percentage of MAP2-positive neurons with at least one perinuclear GA focus without (A) nuclear DSBs within the same cell vs cells with both GA and at least three DSB foci (B). Note the striking concurrence of perinuclear GA and nuclear DSB foci within same neurons at D80 in parental C9 and C9-KO as opposed to nearly absent foci of either type in all other conditions. **(b, c)** Asterisks: highly significant increase in any pairwise comparison with unlabeled conditions, one-way ANOVA with Bonferroni post test, *P≤0.05, **P≤0.01, ***P≤0.001, N=60 images from 3 independent experiments, error bars=SD. All unlabeled conditions (i.e. with no asterisk) were not significantly different amongst themselves in any pairwise comparison.

**Figure 7.**
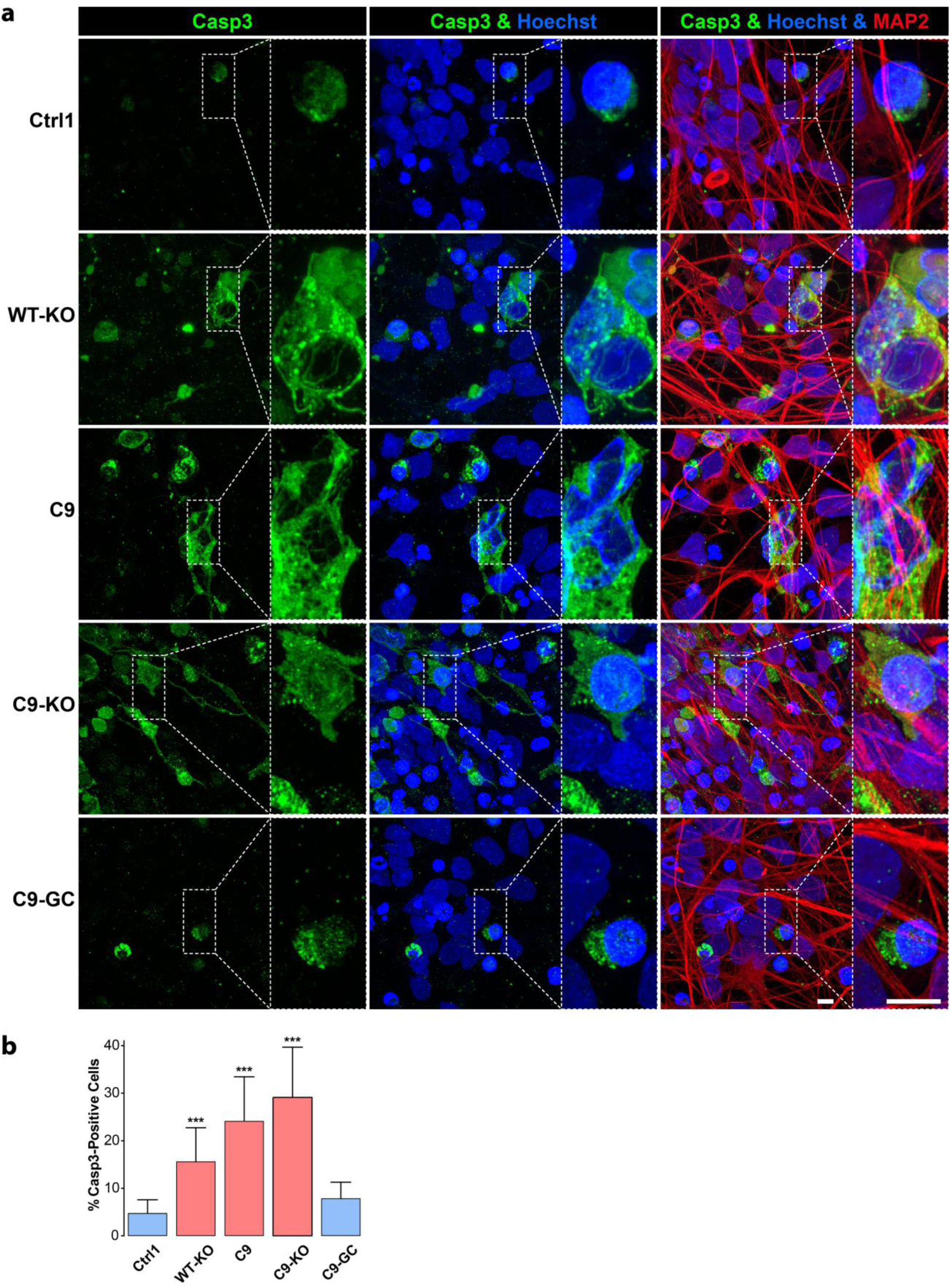
Premature apoptosis occured in C9ORF72 spinal MNs over ageing. **(a)** Apoptosis in MAP2-positive neurons (in red) was revealed as cytosolic rim staining around Hoechst-positive nuclei (in blue) for cleaved caspase 3 (Casp3, in green) by confocal IF microscopy at D80 endpoints (fig. 3). Dotted boxed areas in image galleries are shown magnified on the right. Note the striking accumulation of Casp3 in parental C9 and C9-KO that was phenocopied by WT-KO. Conversely, apoptosis hardly occurred in C9-GC and parental Ctrl1. Scale bars = 10µm. **(b)** Quantification of (a), percentage of apoptotic cells in MAP2-positive population. Asterisks: highly significant increase in any pairwise comparison with unlabeled conditions, one-way ANOVA with Bonferroni post test, *P≤0.05, **P≤0.01, ***P≤0.001, N=60 images from 3 independent experiments, error bars=SD. All unlabeled conditions (i.e. with no asterisk) were not significantly different amongst themselves in any pairwise comparison. Remaining comparisons were: WT-KO vs C9: ***, WT-KO vs C9-KO: ***, C9 vs C9-KO: ***.

Remarkably, KO of exonic C9ORF in control cells with naturally no HREs mimicked DSB accumulation as high as in C9 or C9-KO lines (WT-KO, fig. S6a, b), suggesting that LOF of C9ORF72 is the driving factor for appearance of DSBs rather than augmented DPRs. By contrast, gene correction of HREs in C9ORF reverted augmented GP, GA and DSBs back to control levels (C9-GC, fig. 5, 6), consistent with the axonal trafficking profiles (fig. 3b).

The concurrence of augmented DPR and DSB foci in C9 and C9-KO at D80 (fig. 5, 6) raised the question whether either type of perturbation concurred in the same neuron or in two distinct parts of the population. This presents an important aspect of C9ORF72 pathology as the causative GOF of elevated DPR expression in DNA damage is hotly debated (*26, 27*). As for the neuritic GP foci, we were unable to assign affected neurites to their respective somata in our dense, aged cultures (fig. 5) whereas we always observed augmented nuclear DSBs in the same neuron whenever perinuclear GA foci emerged (fig. 6a, c, green and white arrowheads in a). In essence, ~40% of all C9 and C9-KO neurons displayed perinuclear GA foci together with nuclear DSBs (fig. 6c). In summary, we conclude that DPR accumulation in aged C9ORF spinal MNs is probably always linked to concurrence of DSBs. Besides, DPR accumulation does not seem to be a requirement for DSBs as WT-KO MNs showed DSBs despite having no HREs.

Finally, axonal trafficking phenotypes and accumulated DSBs in C9, C9-KO and WT-KO at D80 were mirrored by elevated Casp3 levels whereas Ctrl1 and C9-GC were showing hardly any sign of apoptosis (fig. 7). Of note, WT-KO MNs exhibited milder Casp3 levels as compared to parental C9 whereas C9-KO displayed even higher levels (fig. 7b), consistent with the smaller/partial imperfect mimic of trafficking defects in WT-KO and more aggravated trafficking defects in C9-KO (fig. 3b).

### TDP43 localization remained unaltered in aged C9ORF72 MNs

Another hallmark in HRE-mediated ALS is the nuclear displacement of TDP43 along with its aggregation in cytosolic inclusion bodies (*36–39*) leading to erratic transcription, splicing, sequestration of RNA and RNA binding proteins and finally deficient RNA granules transport in axons (*40*). Hence, we wished to investigate whether the axonal trafficking defects along with DSB and DPR accumulation at D80 in our C9 lines (fig. 3a, b) were linked to such TDP43 proteinopathies. To this end, we revealed TDP43 localization at D80 by IF immunostainings along with γγH2AX (fig. S7a). We found, again, an increase of DSBs in WT-KO, C9, and C9-KO with no alteration in the prominent nuclear localization of TDP43 as compared to Ctrl and C9-GC MNs (fig. S7a). Specifically, we determined a marginal pool of nuclei devoid of TDP43 that was indistinguishable across all C9 and Ctrl MNs (fig. S7b). Moreover, a minor pool of TDP43 in cytosolic foci was present in all lines but with no change in foci count (Fig. S7c) and total TDP43 amount (i.e. total integral intensity, fig. S7d) across all C9 and Ctrl MNs. In conclusion, the phenotypic perturbations in our C9 cell models were not due to upstream TDP43 mislocalizations.

## Discussion

In this study, we wished to compare HRE C9ORF72 phenotypically and mechanistically against mutant FUS and TDP43, all of which are common genetic aberrations causing ALS (*6, 7*). We sought to combine LOF of C9ORF72 with HRE-mediated GOF in a meaningful manner with no overexpression artifacts to clarify the role of both debated mechanisms (*15, 16, 26, 27*). In contrast to FUS- and TDP43 ALS, proximal in parallel to distal axonal trafficking deficits were the hallmarks of C9ORF72 pathology in hiPSC-derived MNs along with accumulation of DPRs, DNA damage and cell death. While GOF and LOF are both contributing to the trafficking deficiencies, C9ORF72 LOF was sufficient to induce DNA damage accumulation and cell death.

Our method of choice was fast dual channel live imaging of axons in compartmentalized spinal MNs cultures, because impaired microtubule-based organelle transport logistics are particularly vulnerable in these long neurites and often debated as possible cause for neurodegeneration (*8–10*) as they impact on diverse crucial processes such as signal progression, neurotrophic and nutritional support, target finding, neuronal plasticity and regeneration, denervation, energy support, local deposition of mRNA, etc. Moreover, such trafficking defects do not occur independently from other pathomechanistic events (*1*) such as aggregate depositions along with altered nucleo-cytosolic shuttling of disease mediators (*5, 41*), suppression of neuroprotective HSP induction (*42, 43*), impaired DNA damage response and repair (*5, 44, 45*). For example, we have recently shown that mutations in the nuclear location sequence of FUS hampers its nuclear import and causes its cytosolic aggregation along with its failure to recruit the nuclear DNA repair machinery to damage sites (*5*). Remarkably, this impaired DNA damage response had a profound detrimental impact on distal axonal organelle motility and mitochondria activity whereas interfering with microtubule integrity (nocodazole) or oxidative phosphorylation (Oligomycin A) did cause different phenotypes (*33*). Thus, following the paradigm of Modular Cell Biology (*46*) we have further refined our live imaging to obtain multiparametric HC signatures to go beyond a purely descriptive phenotyping of axonal trafficking and gain a predictive systems view of other pathomechanistic pathways as well. We have recently reported distinct signatures for mutant TDP43 and FUS (*33*) (fig. 1b). Given that both proteins are functionally overlapping with roles in DNA/RNA transport (*47*) their distinct signatures demonstrated the power of our multiparametrization to resolve even subtle phenotypic differences.

As for HRE C9ORF72, we found very distinct axonal trafficking defects: (i) the onset of first parameter deviations was at D40 for all lines (fig. 3a) whereas TDP43 and FUS mutants showed severe axonal trafficking deficits as early as D21 (*5, 11, 33*); (ii) distal and proximal axon phenotypes arose simultaneously from D40 onwards whereas all control lines exhibited only distal phenotypes (Fig. 3a, S2); (iii) HRE C9ORF72 signatures were of distinct shape as compared to TDP43 and FUS in fig. 1b with subtle differences between lines presumably due to different genetic backgrounds and HRE repeat numbers (Fig. 3a, S2).

These differences pointed to a distinct spatiotemporal progression of axonal trafficking defects independent of dying-back in HRE C9ORF72 as compared to mutant TDP43 and FUS and were determined by the combinatorial interplay of GOF and LOF mechanisms. As for LOF, KO of the exonic C9ORF72 part in wild type control MNs (WT-KO) led to a partial mimic of the natural HRE C9ORF72 phenotype with a premature onset of distal and delayed proximal trafficking defects (fig. 3b). The same KO in the presence of functional HRE’s (C9-KO) led to the most severe and earliest phenotype, suggesting that both, GOF of HRE and LOF of C9ORF, are driving the axon trafficking deficits. Excision of HREs in C9ORF72 (C9-GC) led to full recovery and control-like signatures (fig. 3b), identifying HREs as the primary trigger of all pathology, i.e. of GOF through RAN-translated DPRs (*26, 27*) along with HRE-mediated LOF through reducing exonic C9ORF72 expression levels.

Next, as HRE pathology is believed to be mediated through RAN-translated DPRs (*26, 27*) we sought to score directly for them when axonal trafficking defects emerged. We succeeded in detecting augmented neuritic GP (fig. 5) and perinuclear GA (fig. 6) foci in parental C9 and C9-KO to similar extends at D80 but not at D21 (fig. S4, S5). As expected, all other lines did not exhibit DPR foci at any time due to no HREs including C9-GC MNs (fig. 5, 6, S4, S5). The concurrence of axonal trafficking defects (fig. 3, S2) with augmented DPRs provides already itself a sufficient explanation of the underlying mechanism as a pending report (under review at Science translational medicine) shows that DPRs can directly inhibit microtubule-based axonal transport of mitochondria and RNA granules by interfering with kinesin-1 and dynein in vivo as well as in vitro-recapitulated (i.e. cell-free) motility assays using recombinant DPRs (*48*). Moreover, we found augmented DSBs at D80 concurring with DPR foci within same MNs in C9 and C9-KO (fig. 5, 6), consistent with recent reports documenting the causative role of DPRs in DSB accumulation (*26, 27*). Furthermore, augmented DPR and DSB foci emerged along with apoptosis (fig. 7), thereby suggesting a causative contribution of HRE-mediated pathology to neurodegeneration as reported (*26, 27*).

However, to our surprise we revealed similar DSB accumulation in WT-KO MNs at D80 (fig. S6b) despite absent HREs along with apoptosis albeit to a milder extend (fig. 7b) and a milder emergence of trafficking defects (fig. 3b). This is of note since yet, all reports pointed towards HRE-mediated DSBs (*26, 27*) and did not accuse LOF of C9ORF as underlying cause for DNA damage accumulation. Furthermore, LOF in wildtype HRE conditions induced a distal axonal phenotype with further aggravation during aging proceeding also to the proximal side (Fig. 3b). This phenotype was reported to be induced by DNA damage induction (*5*) suggesting DNA damage as upstream event in C9ORF72 knockout conditions.

Conversely, parental HRE C9ORF72 exhibited a distinct non-classical dying back phenotype in axons but further exacerbated trafficking defects and apoptosis in C9-KO (fig. 3b, 7b). This was, however, not associated with a further exacerbation in DPR and DSB foci at D80 (fig. 5, 6) but instead even with a 50% reduction in DPR levels in C9-KO cells by ELISA measurements as compared to parental C9, albeit at earlier time points (*32*). However, DSBs were mainly found in DPR containing neurons (Fig. 6c). Again, this finding argues against DPR-mediated DSB accumulation and apoptosis as sole underlying cause in parental HRE C9ORF72. Other GOF mechanisms via detrimental RNA repeat-expansion foci (RREs) and R-loops occurring upstream of DPR translation (*27, 28*) can feasibly explain a certain degree of DPR-independent DSB accumulation. We envision three different explanations for these conflicting data: (i) HRE-mediated GOF contributes to DSB accumulation but is not the sole cause, i.e. similar to the trafficking defects (fig. 3) both GOF and LOF damage DNA concomitantly; (ii) HRE-mediated GOF is the upstream trigger and causes DSB accumulation through the LOF of exonic C9ORF72, thus the KO of C9ORF72 in control cells leads to its LOF in the absence of HREs and consequently to DSB accumulation; (iii) the KO of exonic C9ORF72 in control cells activates the otherwise silent RAN translation of the normal hexanucleotide repeats (nHRs) resulting in DPR expression of distinct composition that escapes detection through available antibodies (GP, GA) as they are too short or other DPR variants accumulate and cause DNA damage. Case (i) and (ii) raise the question of how a LOF of exonic C9ORF72 can cause DSBs. Since C9ORF72 has documented roles in endocytosis with its RNAi-mediated KD impacting on lysosomal degradation and autophagy (*18*), its LOF in ALS could stress cells through perturbed trafficking, protein turnover and ROS accumulation (*49–51*), eventually leading to DSB accumulation as neurons are dying. Case (iii) raises the question of how the KO of exonic C9ORF72 can activate RAN translation of the fewer intronic nHRs in the wild type. Several reports have highlighted the impact of HRE-mediated RAN translation on C9ORF72 expression (*15–17*) but this downregulation is unlikely to be a one-way effect because KO of the exonic C9ORF72 in parental HRE cells reduced vice versa HRE-mediated GP expression (*32*) by 50%, thereby indicating a mutual impact of both the intronic HRE and exonic C9ORF72 part in the RNA transcript on their respective counterpart translation. Since the KO of C9ORF72 was realized by excision of the start codon, an alternative translation start further inwards along with a premature stop codon could feasibly lead to a nonsense-mediated decay of the RNA transcript, thereby limiting HRE-mediated DPR expression whereas in the WT-KO the lack of HREs could instead confer enhanced RNA stability, thereby activating RAN translation of nHRs or at least augmenting RRE foci and R-loops sufficient to increase DSBs (*27*). This explanation appears feasible as overexpression of DPR constructs of quasi wild type length was sufficient to produce RRE foci and R-loops, albeit to a lesser extend (*27*). Consistently, treatment of iPSC-derived control MNs with recombinant DPRs of only 20 repeats were shown (currently under revision at Science translational medicine) to be sufficient to inhibit the microtubule-based motor proteins kinesin-1 and dynein in axons (*48*).

In conclusion, spatiotemporal disease progression of axonal organelle trafficking was distinct in HRE C9ORF72 as compared to FUS and TDP43 (*5, 11, 33*) due to concomitant GOF and LOF mechanisms causing trafficking defects along with DPR and DSB accumulation. While our method of multiparametric HC phenotypic profiling was powerful in revealing these different mutation type-dependent pathomechanisms per se, the detailed dissection of HRE-mediated GOF versus LOF of C9ORF72 in causing DNA damage and neurodegeneration requires further investigations. Many of the reported GOF mechanisms ranging from RRE foci, erratic transcription and splicing (*26, 27*), R-loops (*27, 28*), DPR-mediated DSBs (*26, 27*) and axonal trafficking defects (*48*) are likely to contribute to various extends to the overall pathology whereas the necessity of concomitant LOF highlights a novel aspect calling to investigate the underlying mechanism that presumably comprises the known roles of C9ORF72 in endocytosis and autophagy (*18, 20, 50*). TDP43 did not exhibit any alteration in its prominent nuclear localization even when the most severe trafficking and DSB phenotypes occurred, thereby arguing against an upstream role in the interplay of GOF and LOF mechanisms or otherwise pointing to limitations of our cell model. Nevertheless, our rescue data on C9- GC MNs clearly indicate the therapeutic value of intervening specifically against the HREs in the C9ORF72 locus. Such approaches using antisense oligonucleotides (ASOs) selectively targeting HRE transcripts already exist and yielded promising results (*22*). ASOs are certainly effective in reducing DPR expression and further studies should be performed to clarify if this interference is sufficient for clinical application as other GOF mechanisms stemming from RRE foci and R-loops (*27*) are likely to persist. Moreover, our LOF data (WT-KO) of partial HRE C9ORF72 mimic clearly emphasize the need of further refinement to ensure clinical ASO applications leave C9ORF72 expression unaltered. We propose to further boost the efficacy of ASOs in eliminating HRE-mediated DPR expression by co-targeting RPS25, a small ribosomal protein subunit of 25 kDa required for RAN-translation of DPRs that was recently identified in a genetic screen (*52*).

## Material and methods

### Generation, gene-editing and differentiation of human iPSC cell lines to MNs in MFCs

All procedures were performed in accordance with the Helsinki convention and approved by the Ethical Committee of the Technische Universität Dresden (EK45022009, EK393122012). Patients and controls gave their written informed consent prior to skin biopsy. The generation and expansion of iPSC lines from healthy control and familiar ALS patients with defined mutations in the FUS or TDP43 gene and HREs in C9ORF72 (table 1) was recently described (*5, 11, 17, 53–55*). Isogenic C9-KO and C9-GC lines from parental C9 (table 1) were generated by CRISPR-Cas9n-mediated gene-editing and fully characterized in the Sterneckert laboratory (*32*). The subsequent differentiation to neuronal progenitor cells (NPC) and further maturation to spinal motor neurons (MN) was performed as described (*5, 56*). The coating and assembly of MFCs to prepare for the seeding of MNs was performed as described (*5*). MNs were eventually seeded for final maturation into one site (fig. 1a) of MFCs to obtain fully compartmentalized cultures with proximal somas and their dendrites being physically separated from their distal axons as only the latter type of neurite was capable to grow from the proximal seeding site through a microgroove barrier of 900-µm-long microchannels to the distal site. All subsequent imaging in MFCs (fig. 1a) was performed at D14, 21, 28, 40, 50, 60, 80 of axon growth and MN maturation (D0 = day of seeding into MFCs).

### Live imaging of MN in MFCs

Movie acquisition at strictly standardized readout windows at the distal exit and the proximal entry of the MFC microchannels (fig. 1a) was performed as described (*5, 33*). To track lysosomes and mitochondria, cells were double-stained with live cell dyes Lysotracker Red DND-99 (Molecular Probes Cat. No. L-7528) and Mitotracker Deep Red FM (Molecular Probes Cat. No. M22426) at 50nM each. Trackers were added directly to culture supernatants and incubated for 1 h at 37°C. Live imaging was then performed without further washing of cells in the Center for Molecular and Cellular Bioengineering, Technische Universität Dresden (CMCB) light microscopy facility on a Leica HC PL APO 100x 1.46 oil immersion objective on an inversed fluorescent Leica DMI6000 microscope enclosed in an incubator chamber (37°C, 5% CO2, humid air) and fitted with a 12-bit Andor iXON 897 EMCCD camera (512×512 pixel, 16µm/pixels on chip, 229.55 nm/pixel at 100x magnification with intermediate 0.7X demagnification in the optical path through the C-mount adapter connecting the camera with the microscope). For more details, refer to https://www.biodip.de/wiki/Bioz06_-_Leica_AFLX6000_TIRF and our previous publication (*5*). Fast dual colour movies were recorded at 3.3 frames per second (fps) per channel over 2 min (400 frames in total per channel) with 115 ms exposure time as follows: Lysotracker Red (excitation: 561nm Laser line, emission filter TRITC 605/65 nm) and Mitotracker Deep Red (excitation: 633nm Laser line, emission filter Cy5 720/60 nm). Dual channel imaging was achieved sequentially by fast switching between both laser lines and emission filters using a motorized filter wheel to eliminate any crosstalk between both trackers.

Our live set up (at 100x magnification and using the Andor camera as described above) covers in its viewing field at each readout position 2 channels in parallel, each with 117.53µm of their entire length (900µm) from either the distal exit or from the proximal entry.

### Phenotypic HC profiling

We recently published a comprehensive description of the whole automated analytical pipeline starting from object recognition in raw movie data to final multiparametric signature assembly (*33*). In brief, organelle recognition and tracking was performed with the FIJI Track Mate plugin, organelle shape analysis with our custom-tailored FIJI Morphology macro. Both tools returned a set of 9 master parameters in total for each organelle type (mito- versus lysotracker) and readout position (i.e. distal versus proximal), e.g. mean speed and diameter. Subsequent data mining of individual per-movie result files was performed in KNIME to assemble complete final result files with annotated per-organelle parameters, thereby allowing to pool all data of each experimental condition (e.g. all data for mitotracker at the distal readout position at D21 for cell line Ctrl1). In addition, two post-processing parameters were calculated in KNIME, i.e. the Track Ratio Anterograde/Retrograde movement (Fig. 1b, parameter 9) as described (*33*) and the Percentage (of) Moving Tracks (Fig. 1b, parameter 10) as defined as the percentage of tracks with a minimum track displacement of 1.2 µm as arbitrary threshold for moving organelles as opposed to stationary ones. Thereby, a total number of 11 master parameters was finally obtained for each organelle type and readout position. Z-scores were calculated for each parameter (*33*) to express its deviation from pooled control lines at the proximal readout and assembled to whole HC signatures comprising a total of 44 parameters (fig. 1b) as the set of 11 master parameters was applied 4 times (i.e. distal versus proximal and mito- versus lysotracker, fig. 1b). Clustering of whole signatures (fig. 1c, d) was performed with the KNIME node “Hierarchical clustering” as described (*33*).

### Immunofluorescence stainings

For immunofluorescence staining, cells were washed twice with PBS without Ca^2+^/Mg^2+^ (LifeTechnologies) and fixed with 4% PFA in PBS for 10 min at room temperature. PFA was aspirated and cells were washed three times with PBS. Fixed cells were first permeabilized for 10 minutes in 0.2 % Triton X solution and subsequently incubated for 1 hour at RT in blocking solution (1% BSA, 5% donkey serum, 0.3M glycine and 0.02% Triton X in PBS). Following blocking, primary antibodies were diluted in blocking solution and cells were incubated with primary antibody solution overnight at 4°C except for the γH2A.X antibody which was kept for only 2h at room temperature on the fixed material. The following primary antibodies were used: mouse anti-yH2A.X (1:500, Millipore #05- 636), rabbit anti-53BP1 (1:1000, Novusbio NB100-304), chicken anti-MAP2 (1:1000, Abcam ab5392), rabbit anti-Casp3 (1:1000, Cell Signaling #9661), rat anti-GP (1:500, clone 18H8) and mouse anti GA (1:500, clone IAI2) were both generously provided from Dieter Edbauer (*26*). Nuclei were counter stained using Hoechst (LifeTechnologies).

### Image quantification and statistics

For IF microscopy on fixed cells, a minimum of 3 independent experiments based on 3 different differentiation pipelines was always performed. Twenty images per experiment of mature MNs at D21 and D80 after seeding into MFCs were examined and the mean numbers per neuron of foci representing DSBs, DPRs, or TDP43 in MAP2-positive masks were determined using the particle analyser of FIJI after thresholding with the Triangle background subtraction algorithm. Casp3-positive cells and TDP43-positive nuclei were counted manually. Statistical analysis was performed using GraphPad Prism version 5.0. If not otherwise stated, one-way ANOVA was used for all experiments with post-hoc Bonferroni post test to determine statistical differences in pairwise comparisons.*P < 0.05, **P < 0.01, ***P < 0.001, ****P < 0.0001 were considered significant. Data values represent mean ± SD unless indicated otherwise.

For HC phenotypic profiling, data of at least 4 independent experiments based on 4 different differentiation pipelines were pooled to calculate Z-scores. We verified that the inter-experimental variability was marginal as compared to the inter-line variability as described (*33*), thereby validating the pooling of data across all experiments.

## Supplementary Materials

Figure S1. Loss of organelle motility in aged C9ORF72 spinal MNs.

Figure S2. HC phenotypic profiling over extended time course revealed global axonal trafficking defects in C9ORF72 MNs over ageing.

Figure S3. Phenotypic strength progressing over time.

Figure S4. Nuclear DSB and neuritic GP foci were not augmented prior to emergence of axonal organelle trafficking defects in C9ORF spinal MNs at D21.

Figure S5. Nuclear DSB and perinuclear GA foci were not augmented prior to emergence of axonal organelle trafficking defects in C9ORF72 spinal MNs at D21.

Figure S6. γH2AX and 53BP1 colocalized and both revealed augmented nuclear DSB foci in aged C9ORF72 spinal MNs.

Figure S7. TDP43 localization remained unaltered in aged C9ORF72 spinal MNs. Movie 1. Mitotracker

Movie 2. Lysotracker

## Supporting information

Movie 1 - Mitotracker

Movie 2 - Lysotracker

Supplementary Section

## Acknowledgements

we acknowledge the great help in cell culture by Anett Böhme, Sylvia Kanzler, Andrea Kempe and Katja Zoschke. The Light Microscopy Facility (LMF) of CMCB (Center for Molecular and Cellular Bioengineering, Technische Universität Dresden) provided excellent support for all live imaging experiments. We thank Ronny Sczech for having programmed the original FIJI/KNIME analytical HC organelle trafficking pipeline. We thank Dieter Edbauer for generously providing GA and GP antibodies.

## Author contributions

A.P., B.K., H.G., M.A.-R., T.M.B, J.S. and A.H. designed all of the experiments. N.K. generated TDP43 lines, J.J. generated isogenic FUS and Ctrl lines, M.A.-R. generated isogenic C9-KO and C9-GC lines from parental C9 and assisted their cell culture. B.K., M.N. and N.K. performed all live cell imaging, B.K. performed some IF microscopy, A.P. performed all phenotypic profiling, most IF microscopy and all image analysis. H. G. further refined the original FIJI/KNIME analytical HC organelle tracking pipeline, helped with the HC tracking and performed the cluster analysis in KNIME. A.H. supervised the project, A.P. and A.H. wrote the manuscript and all other authors critically revised the manuscript.

## Data and materials availability

all data and materials are available upon request to A.H. All data not included in the main manuscript or Supplementary Materials can be provided through deposition on a public server to be determined with the editor.

## References

1. J. D. Rothstein, Current hypotheses for the underlying biology of amyotrophic lateral sclerosis. Annals of neurology 65 Suppl 1, S3–9 (2009).

2. M. Dadon-Nachum, E. Melamed, D. Offen, The “dying-back” phenomenon of motor neurons in ALS. Journal of molecular neuroscience: MN 43, 470–477 (2011).

3. L. R. Fischer et al., Amyotrophic lateral sclerosis is a distal axonopathy: evidence in mice and man. Experimental neurology 185, 232–240 (2004).

4. D. Frey et al., Early and selective loss of neuromuscular synapse subtypes with low sprouting competence in motoneuron diseases. The Journal of neuroscience: the official journal of the Society for Neuroscience 20, 2534–2542 (2000).

5. M. Naumann et al., Impaired DNA damage response signaling by FUS-NLS mutations leads to neurodegeneration and FUS aggregate formation. Nat Commun 9, 335 (2018).

6. R. Chia, A. Chio, B. J. Traynor, Novel genes associated with amyotrophic lateral sclerosis: diagnostic and clinical implications. The Lancet. Neurology 17, 94–102 (2018).

7. H. P. Nguyen, C. Van Broeckhoven, J. van der Zee, ALS Genes in the Genomic Era and their Implications for FTD. Trends in genetics: TIG 34, 404–423 (2018).

8. S. Salinas, L. G. Bilsland, G. Schiavo, Molecular landmarks along the axonal route: axonal transport in health and disease. Curr Opin Cell Biol 20, 445–453 (2008).

9. M. P. Sheetz, K. K. Pfister, J. C. Bulinski, C. W. Cotman, Mechanisms of trafficking in axons and dendrites: implications for development and neurodegeneration. Progress in neurobiology 55, 577–594 (1998).

10. S. Veleri, P. Punnakkal, G. L. Dunbar, P. Maiti, Molecular Insights into the Roles of Rab Proteins in Intracellular Dynamics and Neurodegenerative Diseases. Neuromolecular Med 20, 18–36 (2018).

11. N. Kreiter et al., Age-dependent neurodegeneration and organelle transport deficiencies in mutant TDP43 patient-derived neurons are independent of TDP43 aggregation. Neurobiol Dis 115, 167–181 (2018).

12. E. Majounie et al., Frequency of the C9orf72 hexanucleotide repeat expansion in patients with amyotrophic lateral sclerosis and frontotemporal dementia: a cross-sectional study. The Lancet. Neurology 11, 323–330 (2012).

13. E. Suh et al., Semi-automated quantification of C9orf72 expansion size reveals inverse correlation between hexanucleotide repeat number and disease duration in frontotemporal degeneration. Acta neuropathologica 130, 363–372 (2015).

14. S. Van Mossevelde, J. van der Zee, M. Cruts, C. Van Broeckhoven, Relationship between C9orf72 repeat size and clinical phenotype. Current opinion in genetics & development 44, 117–124 (2017).

15. P. Frick et al., Novel antibodies reveal presynaptic localization of C9orf72 protein and reduced protein levels in C9orf72 mutation carriers. Acta neuropathologica communications 6, 72 (2018).

16. A. J. Waite et al., Reduced C9orf72 protein levels in frontal cortex of amyotrophic lateral sclerosis and frontotemporal degeneration brain with the C9ORF72 hexanucleotide repeat expansion. Neurobiology of aging 35, 1779.e1775–1779.e1713 (2014).

17. R. Sivadasan et al., C9ORF72 interaction with cofilin modulates actin dynamics in motor neurons. Nature neuroscience 19, 1610–1618 (2016).

18. M. A. Farg et al., C9ORF72, implicated in amytrophic lateral sclerosis and frontotemporal dementia, regulates endosomal trafficking. Human molecular genetics 23, 3579–3595 (2014).

19. T. F. Gendron, L. Petrucelli, Disease Mechanisms of C9ORF72 Repeat Expansions. Cold Spring Harbor perspectives in medicine 8, (2018).

20. Y. Shi et al., Haploinsufficiency leads to neurodegeneration in C9ORF72 ALS/FTD human induced motor neurons. Nature medicine 24, 313–325 (2018).

21. A. Hruscha et al., Efficient CRISPR/Cas9 genome editing with low off-target effects in zebrafish. Development (Cambridge, England) 140, 4982–4987 (2013).

22. J. Jiang et al., Gain of Toxicity from ALS/FTD-Linked Repeat Expansions in C9ORF72 Is Alleviated by Antisense Oligonucleotides Targeting GGG GCC-Containing RNAs. Neuron 90, 535–550 (2016).

23. J. G. O’Rourke et al., C9orf72 is required for proper macrophage and microglial function in mice. Science (New York, N.Y.) 351, 1324–1329 (2016).

24. T. Zu et al., Non-ATG-initiated translation directed by microsatellite expansions. Proceedings of the National Academy of Sciences of the United States of America 108, 260–265 (2011).

25. J. D. Cleary, L. P. Ranum, Repeat-associated non-ATG (RAN) translation in neurological disease. Human molecular genetics 22, R45–51 (2013).

26. Y. Nihei et al., Poly-glycine-alanine exacerbates C9orf72 repeat expansion-mediated DNA damage via sequestration of phosphorylated ATM and loss of nuclear hnRNPA3. Acta neuropathologica 139, 99–118 (2020).

27. C. Walker et al., C9orf72 expansion disrupts ATM-mediated chromosomal break repair. Nature neuroscience 20, 1225–1235 (2017).

28. M. Groh, N. Gromak, Out of balance: R-loops in human disease. PLoS genetics 10, e1004630 (2014).

29. J. Chew et al., Neurodegeneration. C9ORF72 repeat expansions in mice cause TDP-43 pathology, neuronal loss, and behavioral deficits. Science (New York, N.Y.) 348, 1151–1154 (2015).

30. J. G. O’Rourke et al., C9orf72 BAC Transgenic Mice Display Typical Pathologic Features of ALS/FTD. Neuron 88, 892–901 (2015).

31. Z. Hao et al., Motor dysfunction and neurodegeneration in a C9orf72 mouse line expressing poly-PR. Nat Commun 10, 2906 (2019).

32. M. Abo-Rady et al., Knocking out C9ORF72 Exacerbates Axonal Trafficking Defects Associated with Hexanucleotide Repeat Expansion and Reduces Levels of Heat Shock Proteins. Stem Cell Reports 14, 390–405 (2020).

33. A. Pal et al., High content organelle trafficking enables disease state profiling as powerful tool for disease modelling. Scientific data 5, 180241 (2018).

34. S. Millecamps et al., Phenotype difference between ALS patients with expanded repeats in C9ORF72 and patients with mutations in other ALS-related genes. Journal of medical genetics 49, 258–263 (2012).

35. R. Madabhushi, L. Pan, L. H. Tsai, DNA damage and its links to neurodegeneration. Neuron 83, 266–282 (2014).

36. M. DeJesus-Hernandez et al., Expanded GGGGCC hexanucleotide repeat in noncoding region of C9ORF72 causes chromosome 9p-linked FTD and ALS. Neuron 72, 245–256 (2011).

37. E. B. Lee, V. M. Lee, J. Q. Trojanowski, Gains or losses: molecular mechanisms of TDP43-mediated neurodegeneration. Nature reviews. Neuroscience 13, 38–50 (2011).

38. M. Neumann et al., Ubiquitinated TDP-43 in frontotemporal lobar degeneration and amyotrophic lateral sclerosis. Science (New York, N.Y.) 314, 130–133 (2006).

39. E. L. Scotter, H. J. Chen, C. E. Shaw, TDP-43 Proteinopathy and ALS: Insights into Disease Mechanisms and Therapeutic Targets. Neurotherapeutics: the journal of the American Society for Experimental NeuroTherapeutics 12, 352–363 (2015).

40. A. Jovicic, A. D. Gitler, TDP-43 in ALS: stay on target...almost there. Neuron 81, 463–465 (2014).

41. D. Dormann et al., Arginine methylation next to the PY-NLS modulates Transportin binding and nuclear import of FUS. The EMBO journal 31, 4258–4275 (2012).

42. M. Tibshirani et al., Dysregulation of chromatin remodelling complexes in amyotrophic lateral sclerosis. Human molecular genetics 26, 4142–4152 (2017).

43. R. Kuta et al., Depending on the stress, histone deacetylase inhibitors act as heat shock protein co-inducers in motor neurons and potentiate arimoclomol, exerting neuroprotection through multiple mechanisms in ALS models. Cell stress & chaperones 25, 173–191 (2020).

44. S. L. Rulten et al., PARP-1 dependent recruitment of the amyotrophic lateral sclerosis-associated protein FUS/TLS to sites of oxidative DNA damage. Nucleic acids research 42, 307–314 (2014).

45. W. Y. Wang et al., Interaction of FUS and HDAC1 regulates DNA damage response and repair in neurons. Nature neuroscience 16, 1383–1391 (2013).

46. L. H. Hartwell, J. J. Hopfield, S. Leibler, A. W. Murray, From molecular to modular cell biology. Nature 402, C47–52 (1999).

47. I. R. Mackenzie, R. Rademakers, M. Neumann, TDP-43 and FUS in amyotrophic lateral sclerosis and frontotemporal dementia. The Lancet. Neurology 9, 995–1007 (2010).

48. L. Fumagalli et al., C9orf72-derived arginine-containing dipeptide repeats associate with axonal transport machinery and impede microtubule-based motility. bioRxiv, 835082 (2019).

49. L. Ferraiuolo, J. Kirby, A. J. Grierson, M. Sendtner, P. J. Shaw, Molecular pathways of motor neuron injury in amyotrophic lateral sclerosis. Nature reviews. Neurology 7, 616–630 (2011).

50. A. Otomo, L. Pan, S. Hadano, Dysregulation of the autophagy-endolysosomal system in amyotrophic lateral sclerosis and related motor neuron diseases. Neurology research international 2012, 498428 (2012).

51. S. Sasaki, Autophagy in spinal cord motor neurons in sporadic amyotrophic lateral sclerosis. Journal of neuropathology and experimental neurology 70, 349–359 (2011).

52. S. B. Yamada et al., RPS25 is required for efficient RAN translation of C9orf72 and other neurodegenerative disease-associated nucleotide repeats. Nature neuroscience 22, 1383–1388 (2019).

53. A. Catanese et al., Retinoic acid worsens ATG10-dependent autophagy impairment in TBK1- mutant hiPSC-derived motoneurons through SQSTM1/p62 accumulation. Autophagy 15, 1719–1737 (2019).

54. C. J. Donnelly et al., RNA toxicity from the ALS/FTD C9ORF72 expansion is mitigated by antisense intervention. Neuron 80, 415–428 (2013).

55. J. Higelin et al., NEK1 loss-of-function mutation induces DNA damage accumulation in ALS patient-derived motoneurons. Stem cell research 30, 150–162 (2018).

56. P. Reinhardt et al., Derivation and expansion using only small molecules of human neural progenitors for neurodegenerative disease modeling. PloS one 8, e59252 (2013).

