## Supplementary Section for "High content live profiling reveals concomitant gain and loss of function pathomechanisms in C9ORF72 amyotrophic lateral sclerosis"

**This file includes:**

1. figures S1 to S7 with legends
2. captions for movies 1 and 2

**Other Supplemental Materials for this manuscript includes the following:**

MPEG4 movies 1 and 2

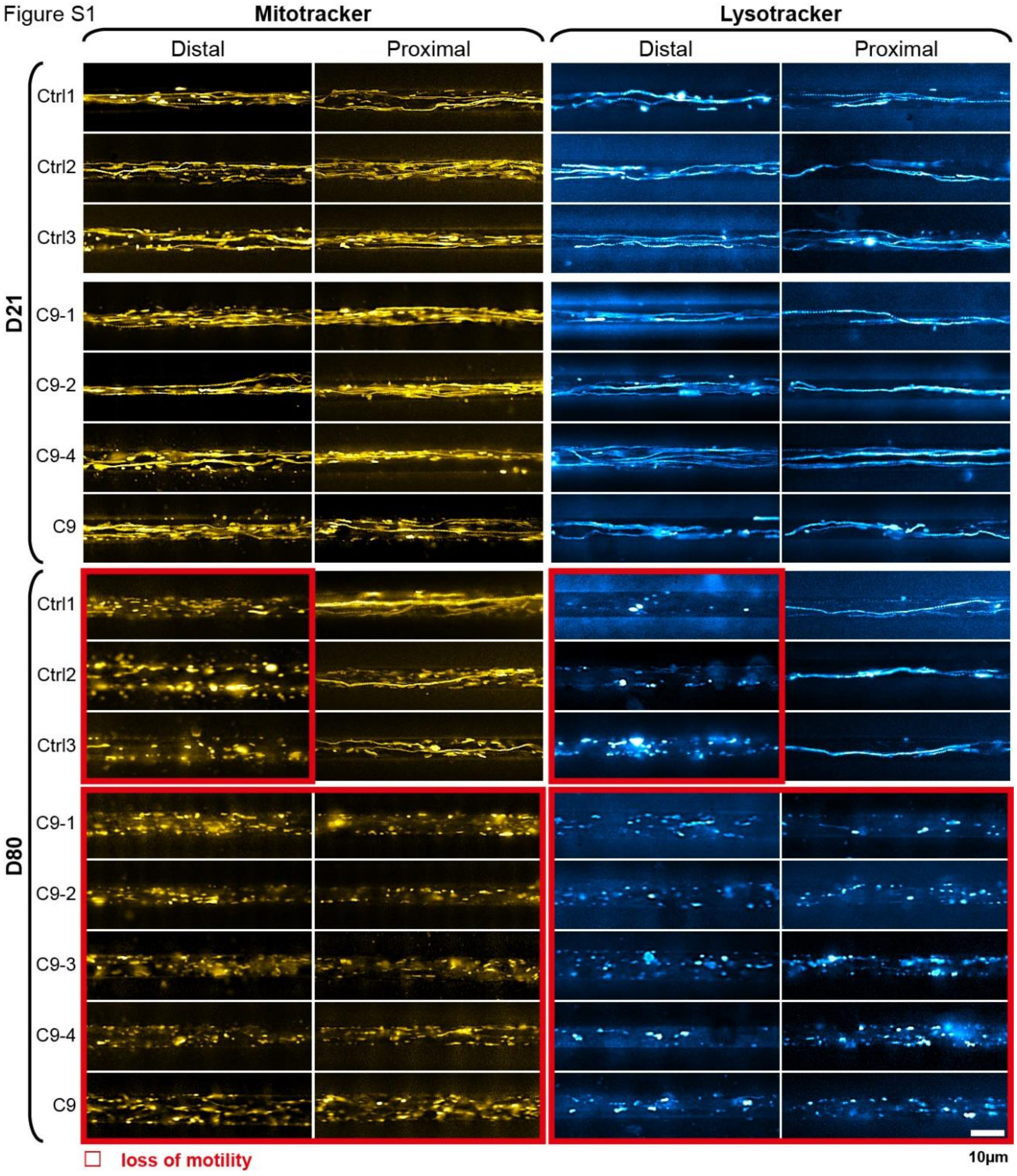


**Figure S1. Loss of organelle motility in aged C9ORF72 spinal MNs.**

Overview of all lines (Ctrls and C9ORFs, table 1), corresponding to fig. 2a. Maximum intensity projections of movie raw data acquired live with Mitotracker (left) and Lysotracker (right) at the distal (left) versus the proximal (right) microchannel readout position as shown in fig. 1a. Movies were acquired at 21 days during maturation (top galleries, D21) versus aged stage at D80 (bottom galleries). Red boxed images highlight loss of motility. Note the loss of motility at D80 simultaneously in both distal and proximal C9ORF axons as opposed to distal loss only in Ctrls. C9-3 was not measured at D21. Scale bar=10µm.

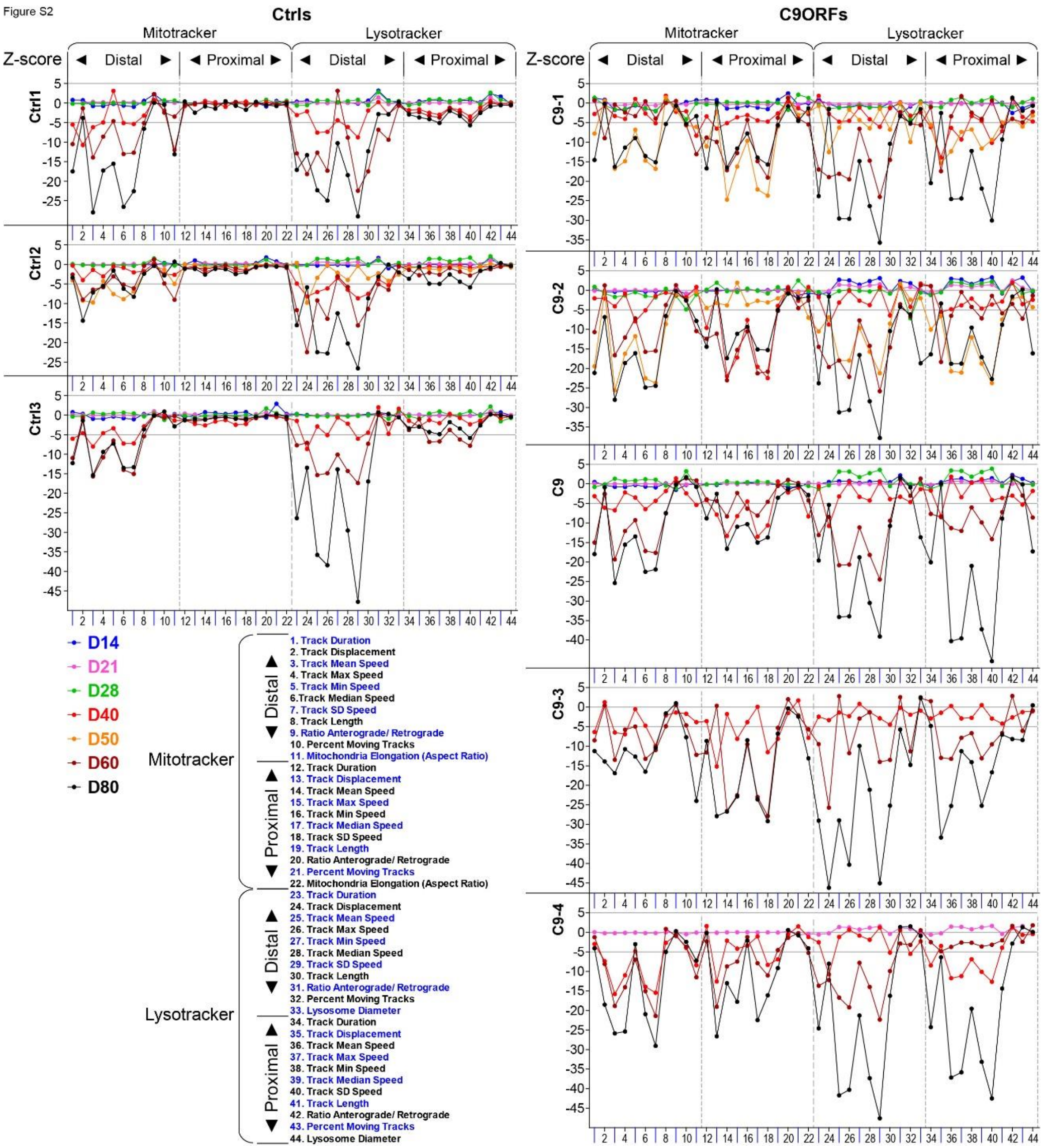


**Figure S2. HC phenotypic profiling over extended time course revealed global axonal trafficking defects in C9ORF72 MNs over ageing.**

Overview of all lines (Ctrls and C9ORFs, table 1), corresponding to fig. 1b and 3a. Multiparametric HC profiles were deduced over a time course for each line from

D14 – D80 (color-coded profiles). C9-3 was not measured at D21, C9-4 was not measured at D50.

Figure S3

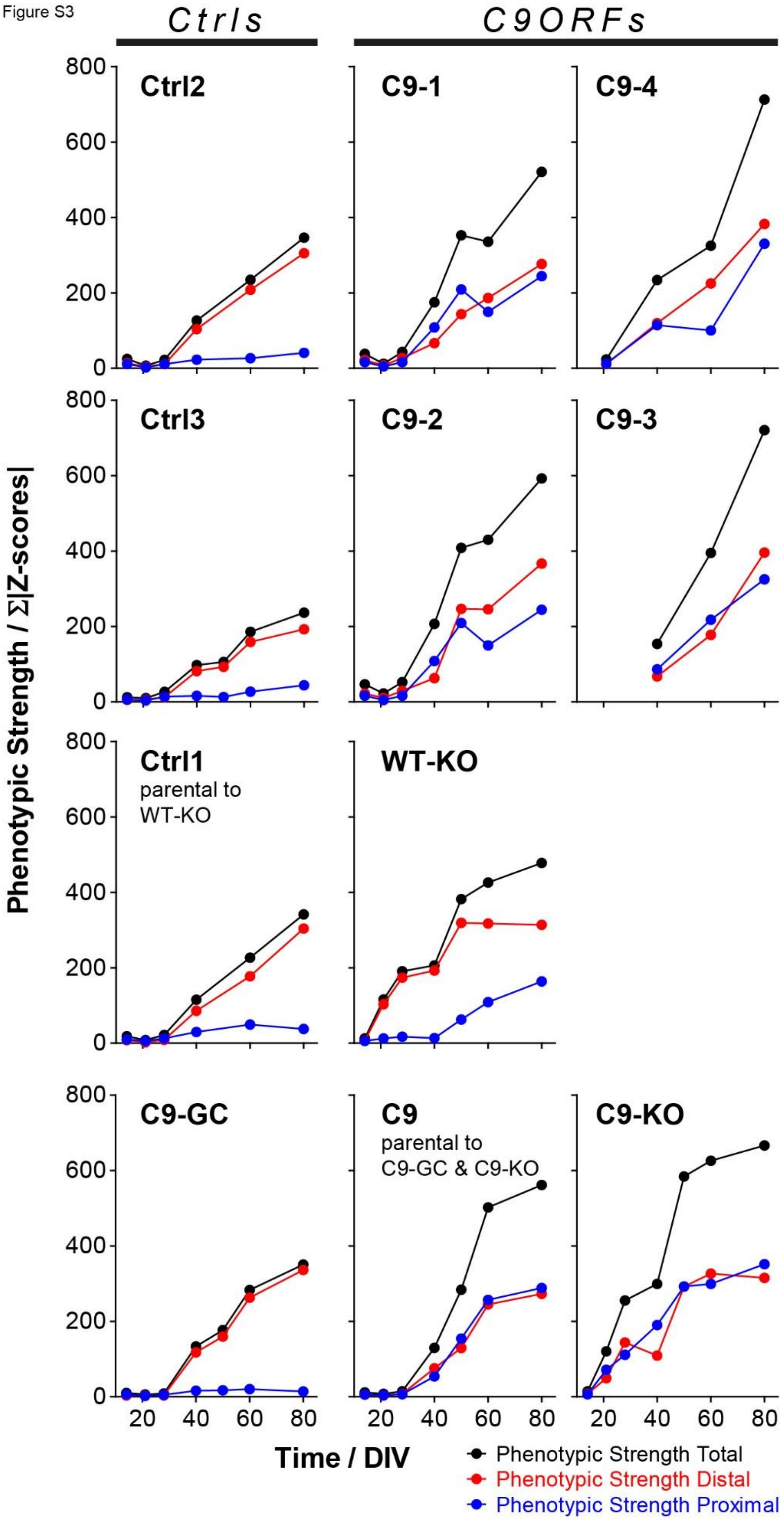

Ctrls
C9ORFs
Ctrl2
C9-1
C9-4
Ctrl3
C9-2
C9-3
Ctrl1
parental to
WT-KO
WT-KO
C9-GC
C9
parental to
C9-GC & C9-KO
C9-KO
Phenotypic Strength / Σ|Z-scores|
0
200
400
600
800
20
40
60
80
Time / DIV
Phenotypic Strength Total
Phenotypic Strength Distal
Phenotypic Strength Proximal

**Figure S3. Phenotypic strength progressing over time.**

Overview of all lines (Ctrls and C9ORFs, table 1), corresponding to fig. 4.

Figure S4

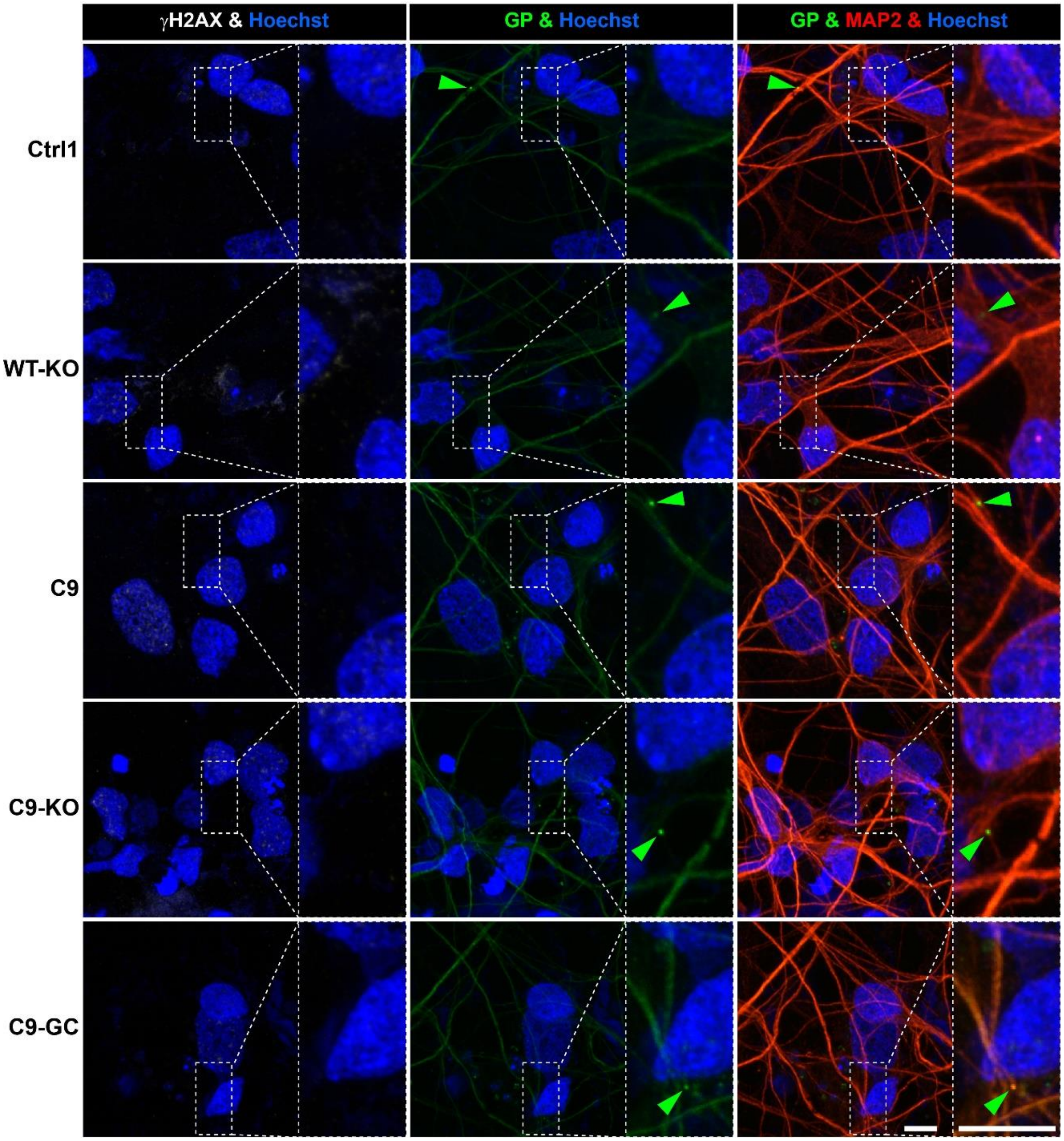


**Figure S4. Nuclear DSB and neuritic GP foci were not augmented prior to emergence of axonal organelle trafficking defects in C9ORF spinal MNs at D21.**

Image gallery at D21, corresponding to D80 in fig. 5. For quantification of nuclear $\gamma$H2AX (in white) and neuritic GP (in green) foci (arrowheads) refer to fig. 5b, c. Scale bars = 10µm.

Figure S5

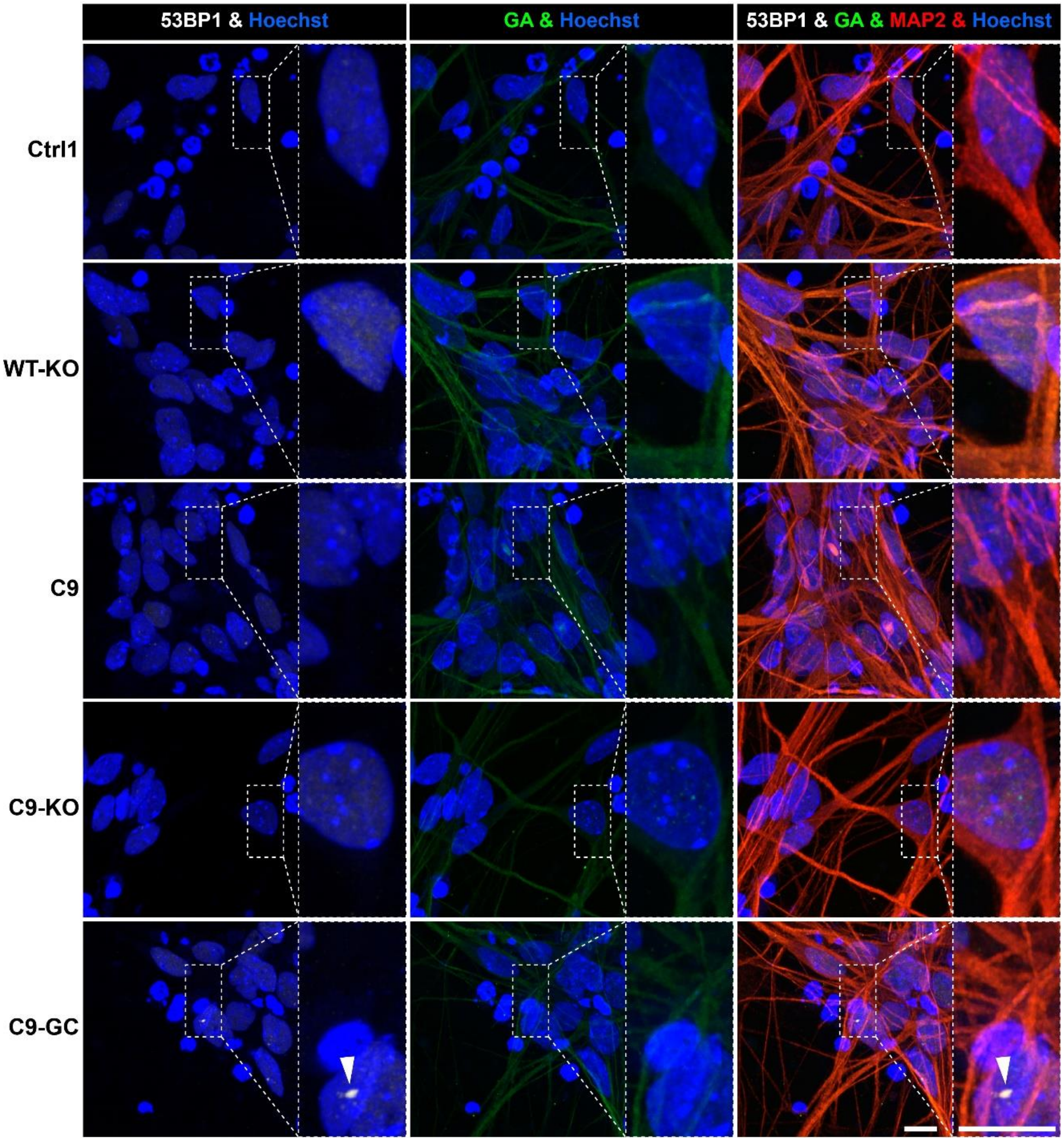


**Figure S5. Nuclear DSB and perinuclear GA foci were not augmented prior to emergence of axonal organelle trafficking defects in C9ORF72 spinal MNs at D21.**

Image gallery at D21, corresponding to D80 in fig. 6. For quantification of nuclear 53BP1 (in white) and perinuclear GA (in green) foci (arrowhead) refer to fig. 6b, c. Scale bars = 10µm.

Figure S6

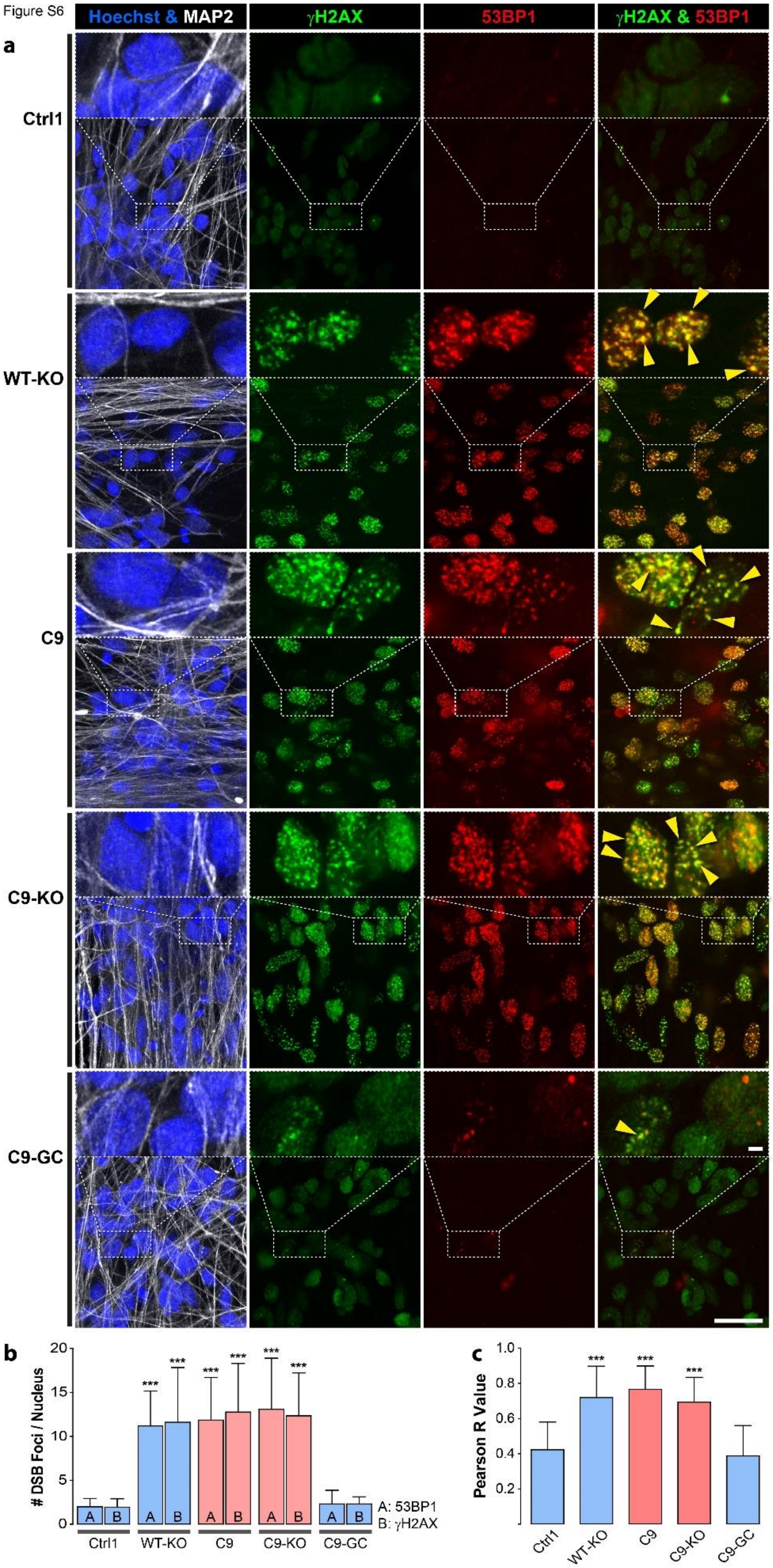
Hoechst & MAP2
γH2AX
53BP1
γH2AX & 53BP1
a
Ctrl1
WT-KO
C9
C9-KO
C9-GC
b
### DSB Foci / Nucleus
20
15
10
5
0
***
A
B
A: 53BP1
B: γH2AX
Ctrl1
WT-KO
C9
C9-KO
C9-GC
c
Pearson R Value
1.0
0.8
0.6
0.4
0.2
Ctrl1
WT-KO
C9
C9-KO
C9-GC

**Figure S6. γH2AX and 53BP1 colocalized and both revealed augmented nuclear DSB foci in aged C9ORF72 spinal MNs.**

**(a)** DSBs were revealed at D80 endpoints by IF staining for γH2AX (in green) and 53BP1 (in yellow) in Hoechst-positive nuclei (in blue) of MAP2-positive (in white) neurons. Note the striking concurrence of both DSB markers in same nuclear foci indicated by yellow overlapping (yellow arrowheads) in parental C9 and C9-KO that was either phenocopied in WT-KO or rescued in C9-GC. Scale bars = 10µm.

**(b)** Quantification (i.e. count) of DSB foci per nucleus in MAP2-positive neurons, left bar (A): 53BP1, right bar (B): γH2AX. Note the similar augmentation of either marker in parental C9 and C9-KO that was either phenocopied in WT-KO or rescued in C9-GC.

**(c)** Quantified colocalization by pixel intensity correlation of both marker channels. Shown are the Pearson correlation coefficients (r values) deduced from the 2D scattergrams by linear regression, r=0: no colocalization, r=1: complete colocalization. Note the high colocalization with mean r~0.7 in parental C9 and C9-KO that was either phenocopied in WT-KO or rescued in C9-GC.

**(b, c)** Asterisks: highly significant increase in any pairwise comparison with unlabeled conditions, one-way ANOVA with Bonferroni post test, $*P \le 0.05$, $**P \le 0.01$, $***P \le 0.001$, N=60 images from 3 independent experiments, error bars=SD. All unlabeled conditions (i.e. with no asterisk) were not significantly different amongst themselves in any pairwise comparison.

Figure S7

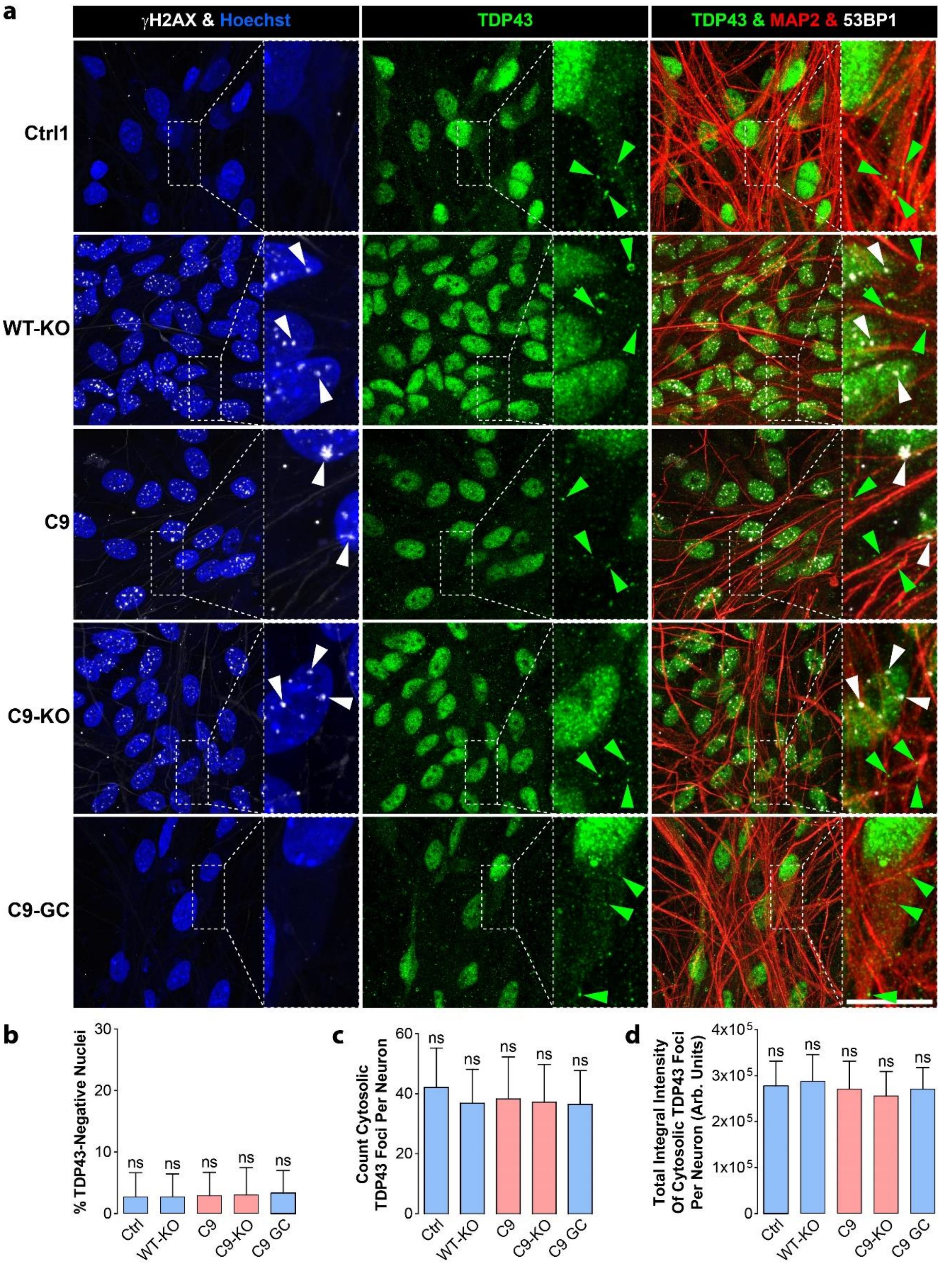
a
γH2AX & Hoechst
TDP43
TDP43 & MAP2 & 53BP1
Ctrl1
WT-KO
C9
C9-KO
C9-GC
b
% TDP43-Negative Nuclei
0
10
20
30
ns
Ctrl
WT-KO
C9
C9-KO
C9 GC
c
Count Cytosolic TDP43 Foci Per Neuron
0
20
40
60
d
Total Integral Intensity Of Cytosolic TDP43 Foci Per Neuron (Arb. Units)
0
1x10^5
2x10^5
3x10^5
4x10^5

**Figure S7. TDP43 localization remained unaltered in aged C9ORF72 spinal MNs.**

**(a)** Nuclear TDP43 (in green) and the DSB marker $\gamma$H2AX (in white) were revealed at D80 endpoints by IF staining in Hoechst-positive nuclei (in blue) of MAP2-positive (in red) neurons. Note the unaltered prominent nuclear TDP43 localization along with unaltered cytosolic foci (green arrowheads) in all Ctrl and C9 lines whereas nuclear DSB foci were increased in WT-KO, C9 and C9-KO (white arrowheads). Scale bars = 10µm.

**(b)** Quantification (i.e. percentage) of TDP43-negative nuclei in MAP2-positive neurons. Note the unaltered nuclear TDP43 localization in all Ctrl and C9 lines.

**(c)** Quantification (i.e. count) of TDP43 cytosolic foci per neuron in MAP2-positive and Hoechst-negative areas. Note the unaltered count in all Ctrl and C9 lines.

**(d)** Quantification of total integral TDP43 intensity of cytosolic foci per neuron in MAP2-positive and Hoechst-negative areas. Note the unaltered amount in all Ctrl and C9 lines.

**(b-d)** ns: no significant change in any pairwise comparison, one-way ANOVA with Bonferroni post test, *$P \leq 0.05$, **$P \leq 0.01$, ***$P \leq 0.001$, N=60 images from 3 independent experiments, error bars=SD.

**Movie 1 and 2 (general remarks)**

All movies were recorded at 100x magnification, NA 1.45 oil immersion, at 3.3 fps per channel over 2 min (400 frames in total per channel) in epifluorescence mode with illumination and filter settings as detailed in Material and methods. All LUTs (Look Up Tables of color-indexed monochromatic raw images) were applied in FIJI software. Organelle motility was revealed with Lysotracker Red DND-99 (LUT Cyan Hot), Mitotracker Deep Red FM (LUT Yellow Hot) at standardized microchannel readout positions distal versus proximal in compartmentalized iPSC-derived MN cultures (fig. 1a), channel length: 900µm.

**Movie 1. Mitotracker**

Refers to fig. 2a (showing maximum intensity projections of Mitotracker Deep Red FM), left, and Lysotracker Red DND-99, right). Shows mitochondria trafficking at D21 versus D80. Note the loss of motility (boxed in red) in C9ORF at D80 at both the distal and proximal readout as opposed to distal loss only in Ctrl cells.

**Movie 2. Lysotracker**

Same as for Movie 1 but for lysosomes.
